# The polyphagous plant pathogenic fungus *Botrytis cinerea* encompasses host-specialized and generalist populations

**DOI:** 10.1101/716985

**Authors:** Alex Mercier, Florence Carpentier, Clémentine Duplaix, Annie Auger, Jean-Marc Pradier, Muriel Viaud, Pierre Gladieux, Anne-Sophie Walker

## Abstract

The host plant is often the main variable explaining population structure in fungal plant pathogens, because specialization contributes to reduce gene flow between populations associated with different hosts. Previous population genetic analysis revealed that French populations of the grey mould pathogen *Botrytis cinerea* were structured by hosts tomato and grapevine, suggesting host specialization in this highly polyphagous pathogen. However, these findings raised questions about the magnitude of this specialization and the possibility of specialization to other hosts. Here we report specialization of *B. cinerea* populations to tomato and grapevine hosts but not to other tested plants. Population genetic analysis revealed two pathogen clusters associated with tomato and grapevine, while the other clusters co-occurred on hydrangea, strawberry and bramble. Measurements of quantitative pathogenicity were consistent with host specialization of populations found on tomato, and to a lesser extent, populations found on grapevine. Pathogen populations from hydrangea and strawberry appeared to be generalist, while populations from bramble may be weakly specialized. Our results suggest that the polyphagous *B. cinerea* is more accurately described as a collection of generalist and specialist individuals in populations. This work opens new perspectives for grey mold management, while suggesting spatial optimization of crop organization within agricultural landscapes.

## Introduction

Fungi are ubiquitous in ecosystems, but most fungal species are not globally distributed and display some kind of population structure (Taylor *et al.*, 2006). In fungal plant pathogens, investigating population structure in relation to host range is particularly relevant, as it is essential for disease management to recognize the different populations of a pathogen, which can have different life history traits, such as resistance to fungicide or phenology (Milgroom & Peever, 2003). Beyond the practical interest in understanding the structure of pathogen populations, studying barriers to gene flow between populations within species may reveal key features of adaptive divergence before they become confounded by other factors. The host plant is often the main variable explaining population subdivision in fungal plant pathogens, because specialization to hosts contributes to reduce gene flow between populations associated with different hosts. Host specialization acts against gene flow due to the reduced viability of migrants and hybrids, and the role of specialization as a barrier to gene flow is reinforced in species that reproduce within their hosts, which is the case for many plant pathogens (Giraud *et al.*, 2010; Giraud *et al.*, 2006; Gladieux *et al.*, 2010). Hence, polyphagous fungal pathogens can form collections of specialist populations, each associated with a limited number of hosts (*i.e.* poly-specialists, such as the rice blast fungus *Pyricularia oryzae*; Gladieux *et al.*, 2018; Valent *et al.*, 2019). Depending on time-based, geographic and genetic factors (Nosil *et al.*, 2009), speciation can then unfold relatively rapidly between pathogen populations (Giraud et al. 2010). The phytopathological literature provides many examples of adaptive radiations (Schluter, 2000) in pathogens, with many sibling pathogen species associated with distinct hosts (Giraud *et al.*, 2006; Tellier *et al.*, 2010). Alternatively, in some pathogens, infection of a novel host is not associated with a specialization process, and merely results in the expansion of the host range of a generalist species. The domestication of plant pathogens, *i.e.* the effective and durable control over pathogen reproduction by humans, remains a major goal in agricultural sciences and knowledge of the intrinsic and extrinsic factors underlying the emergence of new populations specialized onto new hosts –or expansions in host range in generalist species– should help reaching this goal.

*Botrytis cinerea* (Ascomycota) is a textbook example of a highly polyphagous fungal plant pathogen, causing grey mold on more than 1400 known hosts, in 586 plant genera and 152 botanical families, including high-value crops such as grapevine and tomato (polyphagy index = 54; Elad *et al.*, 2016; Supplementary Information 1). *B. cinerea* is a necrotrophic pathogen that asexually produces conidia that form a sporulating grey layer on the surface of infected tissues. Conidia are mainly dispersed during spring and summer by wind and human activity. During winter, the fungus develops as a saprobe on plant debris, where it can proceed to undergo sexual reproduction and form melanin-pigmented sclerotia (Carisse, 2016). The analysis of population structure of *B. cinerea* populations infecting wild (e.g. Giraud *et al.*, 1999; Rajaguru & Shaw, 2010) or cultivated (*e.g.* Karchani-Balma *et al.*, 2008; Leyronas *et al.*, 2015) host plants revealed differentiation between sympatric populations from different hosts. However, no correlation was found between population structure and cross-pathogenicity, suggesting that the observed differentiation between populations was not related to host specialization, or that it was instead linked to components of specialization that have not been measured. For example, Leyronas *et al.* (2015) showed lack of genetic differentiation between *B. cinerea* populations found on tomato (*Solanum lycopersicum*) and lettuce (*Lactuca sativa*). Patterns of pathogenicity were also consistent with generalism, as strains showed significant differences in quantitative pathogenicity on tomato hosts, but not on lettuce. Recently, we have found strong genetic differentiation between isolates collected in the greenhouse on tomato on the one hand and outdoor on bramble (*Rubus fruticosus*) and grapevine (*Vitis vinifera*) on the other hand, suggesting possible specialization of the pathogen to these hosts (Walker *et al.*, 2015). Here, we test the hypothesis of host specialization in *B. cinerea*, by further investigating the genetic structure of *B. cinerea* populations sampled in four French regions on sympatric hosts, including previously tested hosts (*e.g.* grapevine, tomato and bramble; Walker *et al.*, 2015) and new hosts (*e.g.* strawberry-*Fragaria* x *ananassa-* and hydrangea-*Hydrangea macrophylla*), and by measuring quantitative pathogenicity on various hosts using cross-pathogenicity testings. More specifically, we addressed the following questions: (i) What is the population structure of *B. cinerea* on grapevine, tomato, bramble, strawberry and hydrangea in France? (ii) Are there differences in qualitative or quantitative pathogenicity between identified populations? (iii) Are patterns of population structure and pathogenicity consistent with the specialization of *B. cinerea* to some hosts?

## Results

### Population subdivision

We characterized 683 isolates from 5 hosts in 15 sites representing 4 areas of France, using 8 microsatellite markers. To examine population structure, we first characterized the partitioning of genetic variation among different factors (host plant and area) using hierarchical analyses of molecular variance (AMOVA) (**Error! Reference source not found**.). With samples grouped by host, variation among hosts was found highly significant (*P*<0.001; *F*_*ST*_=0.18) and accounted for 18.4% of the molecular variance. Variation due to geography was also found significant but accounted for only 5.5% of molecular variance (*P*<0.001; *F*_*ST*_=0.06). With samples grouped by host and nested within areas, variation within populations accounted for most of the molecular variance (77,7%; *P*<0.001; *F*_*ST*_=0.07). Variation among hosts, and among hosts within areas were significant (*P*<0.001) and accounted for 16% and 6% of the molecular variance, respectively. Variation among areas was not significant (*P*=0.772), but variation among areas within hosts was significant (*P*<0.001) and accounted for 23.1% of the molecular variation, respectively. These results suggest that hosts (tomato, grapevine, strawberry, bramble and/or hydrangea) and areas (Alsace, Champagne, Loire Valley and Provence) explain a significant proportion of the genetic variance in the dataset, with a stronger effect of the host.

We investigated patterns of population subdivision using (i) the Bayesian clustering method implemented in STRUCTURE, assuming a model with admixture and correlated allele frequencies, (ii) the non-parametric multivariate clustering method implemented in the discriminant analysis of principal components (DAPC), and (iii) the maximum likelihood clustering method implemented in SNAPCLUST (Figure 1). The three methods converged towards the same clustering solutions at all *K* value. Models with *K*=5 captured the most salient features of the structure of the dataset, with two clusters mostly associated with grapevine and tomato, respectively, and three clusters found on all hosts but tomato. Increasing *K* above 5 did not reveal new clusters but mostly introduced heterogeneity in clustering patterns (see also Supplementary Information 4 & 5). We calculated the mean pairwise *F*_*st*_ between clusters for each *K* value (Supplementary Information 3) and at *K=5* the *F*_*st*_ is close to its maximum and displays very close values across all three clustering methods. The Δ*K* Evanno method (Supplementary Information 6) favors *K*=4, while goodness of fit statistics (Supplementary Information 7) favors *K*=5. All subsequent analyses assumed *K*=5 and isolates were assigned to clusters using the membership proportions estimated with STRUCTURE. Using membership proportions greater than 0.7 to assign multilocus genotypes, clusters tended to be associated with one main host (Figure 2A). Clusters C1 and C2 were almost exclusively associated with tomato and grapevine, respectively (96% of isolates from C1 were collected on tomato; 84% of isolates from C2 collected on grapevine). The three other clusters were less specific, with the majority of C3 isolates collected on grapevine (53%), and the majority of C4 and C5 isolates collected on bramble (50 and 60%, respectively). Isolates from strawberry were mostly distributed in C3 and C4, whereas isolates collected from hydrangea were mostly assigned to C4 and C5. Pairwise Jost’s *D* between clusters inferred with STRUCTURE were all significantly different from zero and ranged between 0.320 and 0.869 at *K*=5 (Table 3). Highest values were observed for pairs including tomato-associated cluster C1 (range 0.663-0.869) and bramble-associated cluster C5 (range 0.658-0.869). The smallest differentiation was observed between grapevine-associated clusters C2 and C3 (D=0.320).

**Figure 1.**
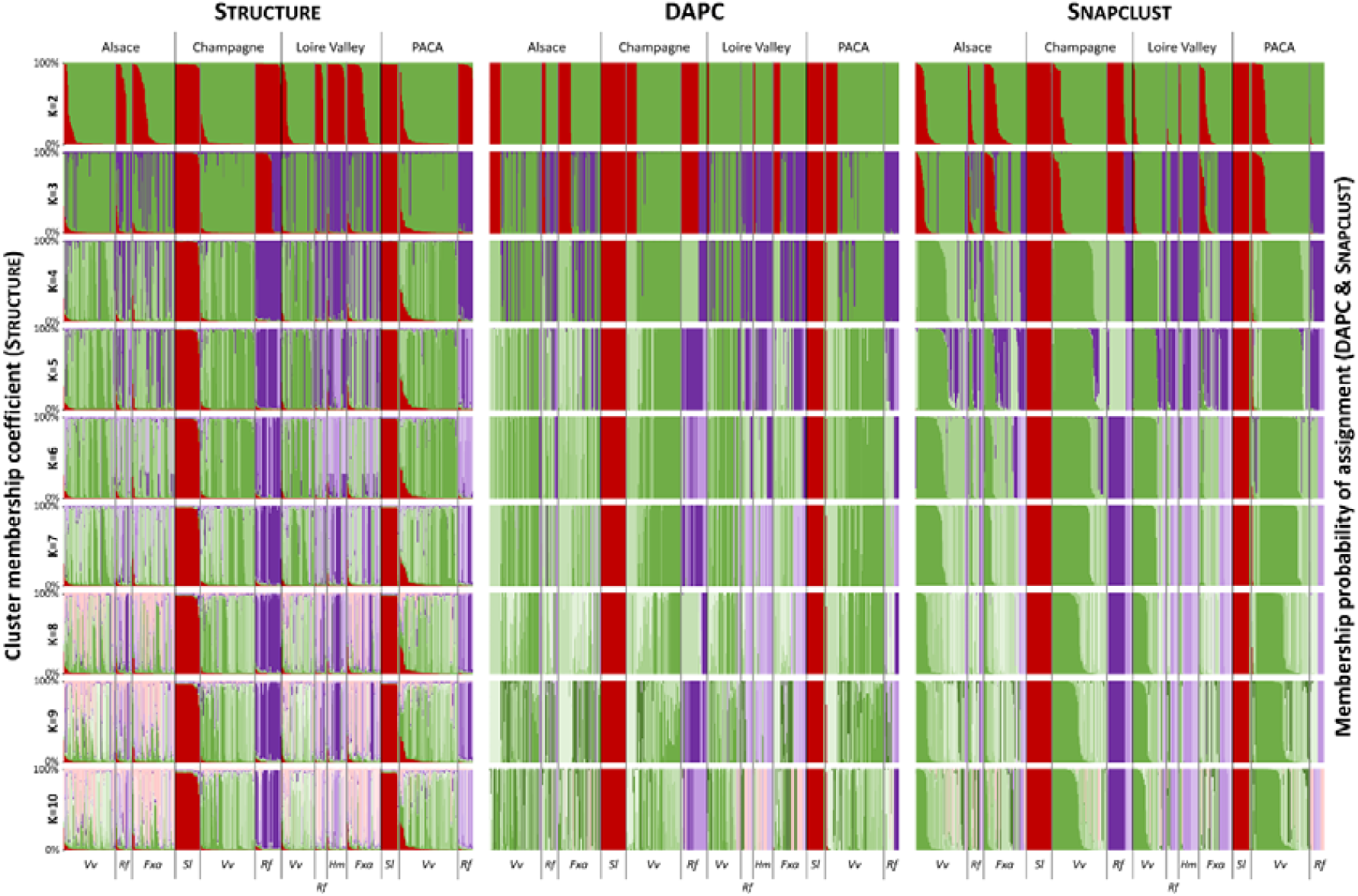
Barplots of the Structure, DAPC and Snapclust analysis showing genetic subdivision of the 681 *B. cinerea* isolates collected between 2002 and 2015 into a varying number of genetic clusters (K=2 to K=10). Individuals are sorted out by their area of origin (black vertical bars) and then by their sampling host (grey vertical bars). The areas names are indicated at the top of the chart and the host of origin at the bottom in abbreviated form (*Vv: V. vinifera; Rf: R. fruticosus; Fxa: F. x ananassa; Sl: S. lycopersicum and Hm; H. macrophylla*). Each cluster is colored according to the most represented host of its members (red for the tomato-associated cluster, shades of green for grapevine-associated clusters, shades of purple for bramble-associated clusters and shades of pink for strawberry-associated clusters).

**Figure 2.**
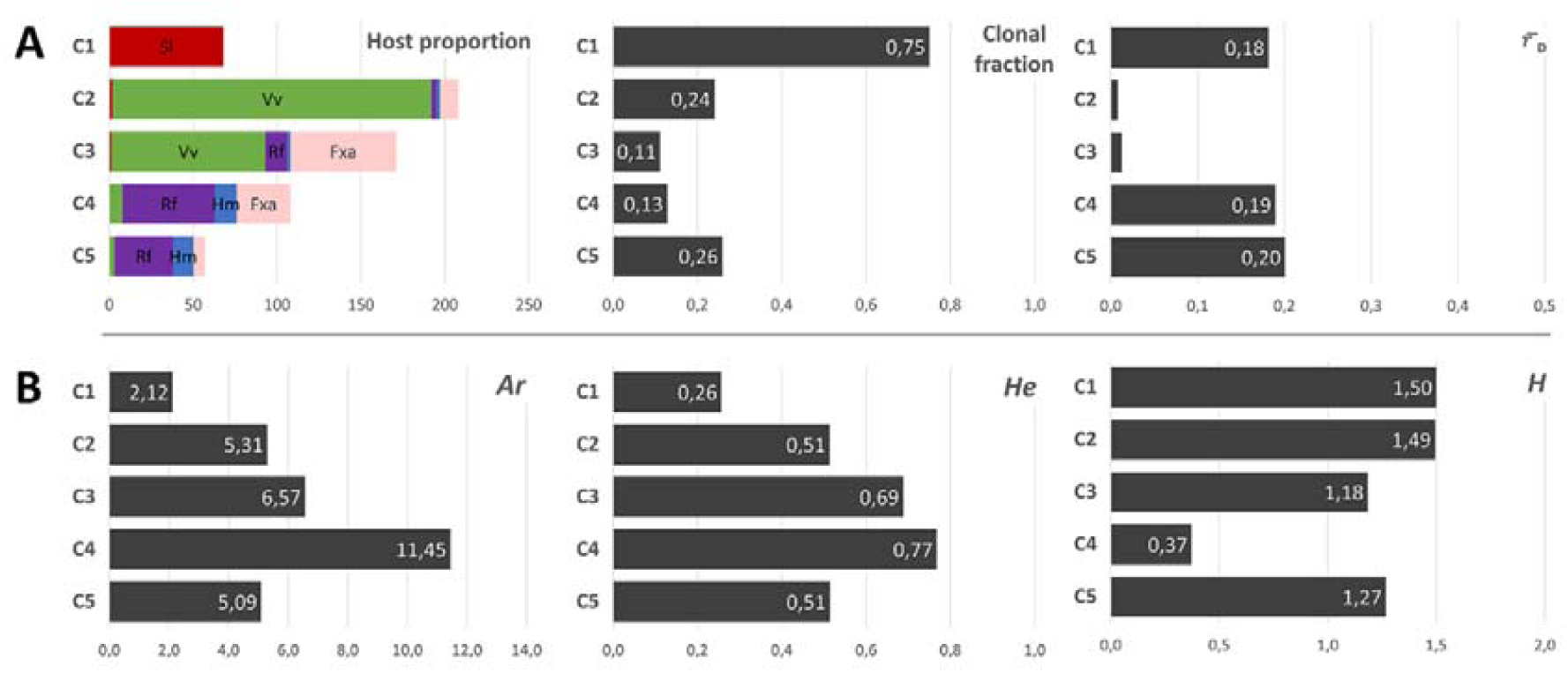
Patterns of diversity and reproduction mode measured in the five clusters delimited by the STRUCTURE partitioning analysis at K=5. (A) Proportions of sampling hosts represented in inferred clusters (left), clonal fraction (middle) and 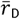 (estimation of multilocus linkage disequilibrium; right). Isolates from tomato Sl are colored in red, grapevine Vv in green, bramble Rf in purple, hydrangea Hm in blue and strawberry Fxa in pink. (B) Diversity is estimated *via* allele richness *A*_*r*_ (mean number of alleles per locus; left), the expected heterozygosity *He* (mean expected heterozygosity over the eight loci; middle) and the Shannon’s diversity index *H* (mean over the eight loci; right).

### Patterns of genetic diversity and reproductive mode

We investigated how genetic variability varied across the clusters defined at *K*=5, by estimating the mean number of alleles per locus *Ar,* genic diversity *He*, and Shannon’s diversity index *H* (Figure 2B). The tomato*-*associated cluster C1 was the least variable (*Ar*=2.12; *He*=0.26), while the bramble-associated cluster C4 was the most variable (*Ar* =11.45; *He*=0.77). Intermediate levels of variability were observed for the grapevine-associated clusters C2 and C3 (Ar=5.31 and 6.57; *He*=0.51 and 0.69) and the other bramble-associated cluster C5 (*Ar*=4.01; *He*=0.44). Multilocus genotypes were evenly represented in clusters C1, C2, C3 and C5 (1.18<*H*<1.5), but not in C4 (*H*=0.37).

To investigate the reproductive mode of *B. cinerea*, we computed the proportion of genotypes repeated multiple times (clonal fraction 1-G/N) and we estimated multilocus linkage disequilibrium (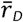; Figure 2A). The highest clonal fraction was found in the tomato*-*associated cluster C1 (CF=0.75), which also displayed a relatively high level of linkage disequilibrium among markers 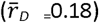. Clonal fraction was low and in the same order of magnitude (range: 0.11-0.26) across other clusters. 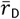 values were also low for these clusters (range: 0.008-0.20), although significantly different from zero.

### Cross-pathogenicity tests

#### On leaves

We assessed differences in aggressiveness between clusters using cross-pathogenicity tests on strawberry, grapevine, bramble, tomato and hydrangea leaves, with bean used as a naive host (no isolates were sampled on this host). Mean necrosis diameters are reported in Figure 3 and differences between pairs of hosts of origin are presented in Table 3. Results of cross-pathogenicity leaf tests observed over post-inoculation time for *B. cinerea* isolates representative of their host of collection.

**Figure 3.**
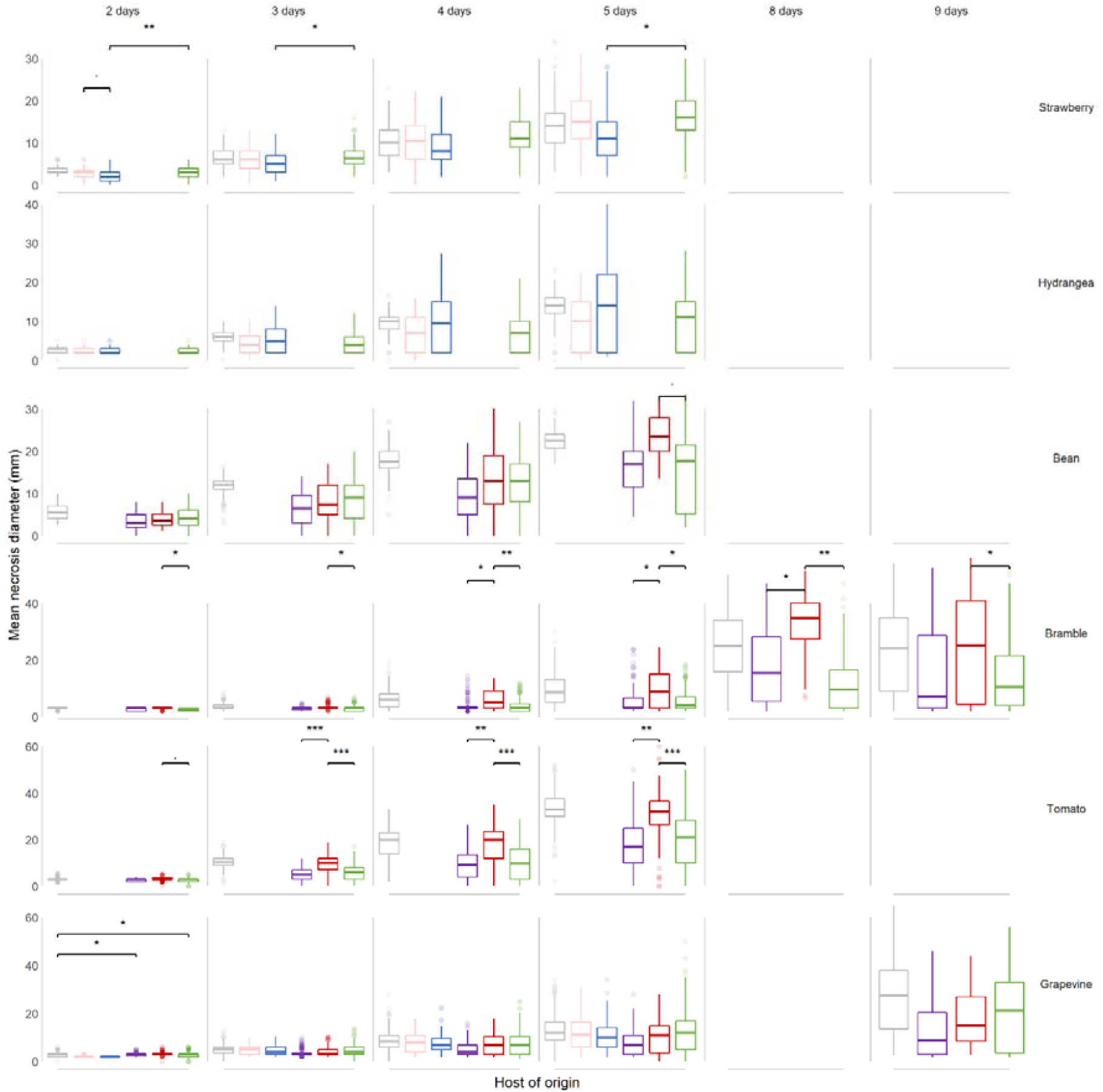
Evolution of mean necrosis diameter over time, for *B. cinerea* isolates representative of their host of origin and inoculated on a range of test hosts. Each boxplot represents a group of 7-33 isolates collected on strawberry (pink), hydrangea (blue), bramble (purple), tomato (red), grapevine (green) and a reference isolate used as a control (grey). These isolate subgroups were tested on leaves from strawberry, hydrangea, bean, bramble, tomato and grapevine (test hosts). Lesions were scored between 2 and 9 days after inoculation, depending on the test host. ., *, ** and *** indicates tests of difference of mean necrosis diameter between two groups of isolates with *P*-value lower than the 10%, 5%, 1% and 1‰, respectively.

**Figure 4.**
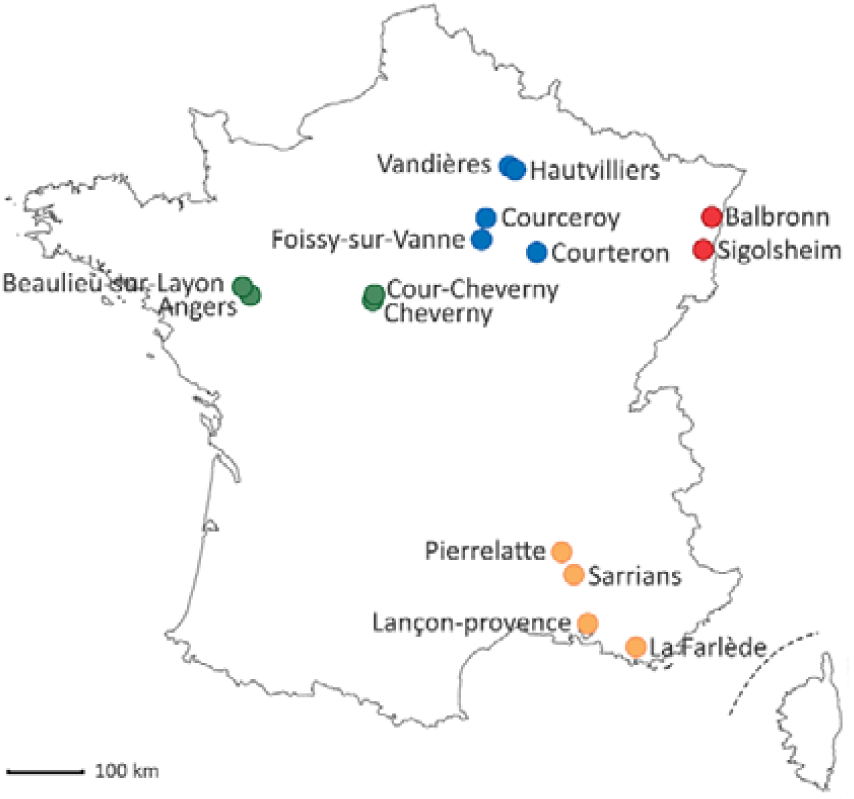
Map of *B. cinerea* populations collected from four French areas on various host plants between 2002 and 2015. Each color represents an area with Alsace in red, Champagne in blue, Loire Valley in green and Provence in yellow.

**Figure 5.**
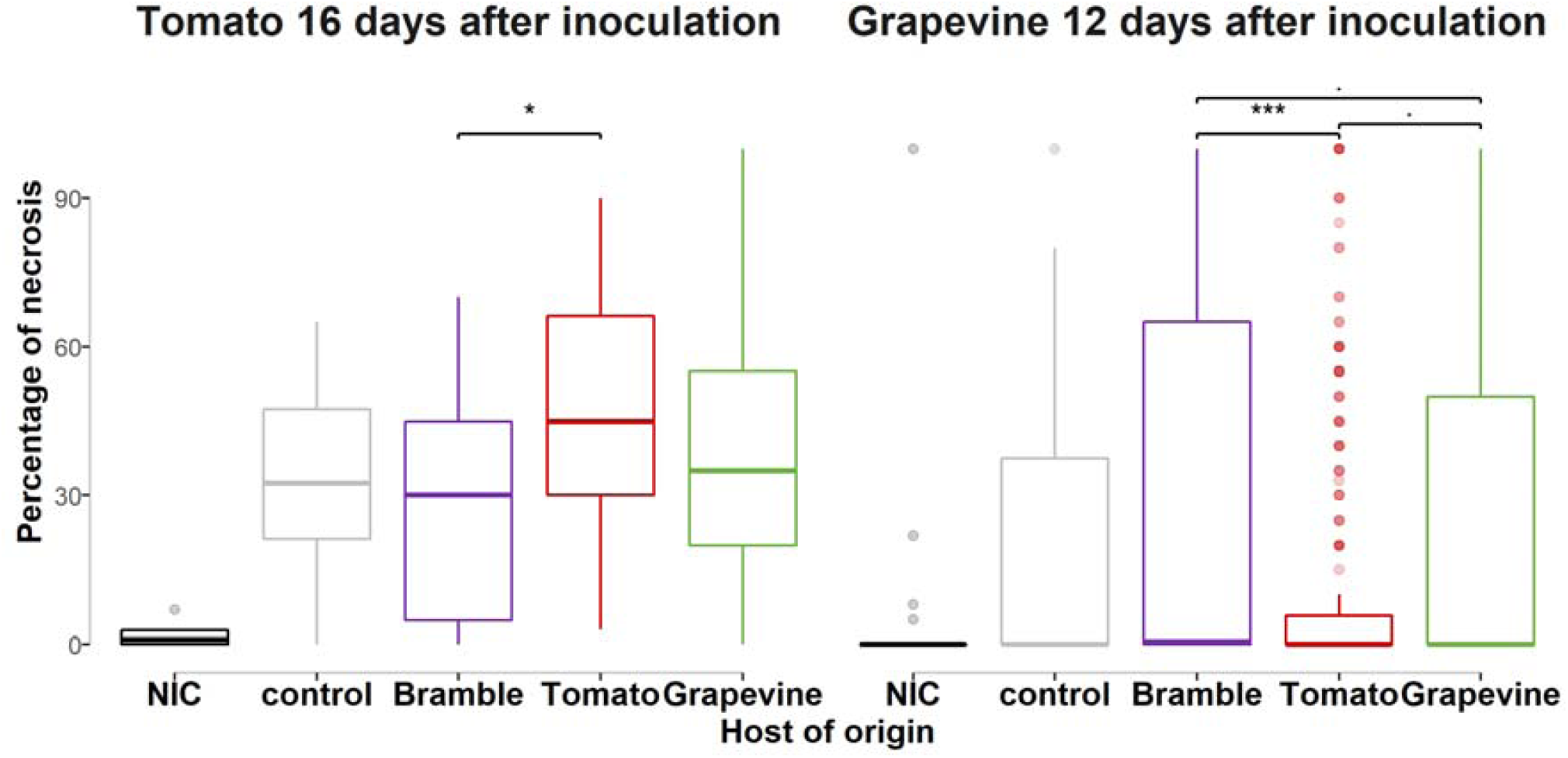
Percentage of necrosis for *B. cinerea* isolates representative of their host of origin and inoculated on grapevine and tomato hosts. Each boxplot represents a group of 16-34 isolates collected on bramble (purple), tomato (red), grapevine (green), a reference isolate used as a control (grey) and a not inoculated control (NIC, black). These isolate subgroups were tested on tomato and grapevine berries (test hosts) and percentage of necrosis were measured 16 days and 12 days, respectively, after inoculation. ., *, ** and *** indicates tests of difference necrosis percentage between two groups of isolates with *P*-value lower than 10%, 5%, 1% and 1‰, respectively.

Isolates collected on strawberry, hydrangea, grapevine, bramble and tomato (hosts of origins, in columns) and a reference isolate used as a control were inoculated on strawberry, hydrangea, bean, bramble, tomato and/or grapevine leaves (test hosts, in row). Necrosis were scored between 2 and 16 days after inoculation, depending on the test host.

Our statistical model detected significant effects of the host of origin on the mean necrosis diameters, for several time points after inoculation. Mean necrosis diameters increased over time for all interactions and clusters, and especially for the reference isolate B05.10, indicating that tests were properly carried out. For a given scoring time, aggressiveness of the reference isolate differed among test hosts, bean and tomato showing faster growth of necrosis compared to other hosts. No significant differences (*P-*values > 5%) in aggressiveness were observed among isolates from different clusters inoculated onto the naive host bean, at any scoring time.

On strawberry leaves, the aggressiveness of isolates collected on hydrangea was significantly reduced (range: −0.75 - −4.47mm) compared to isolates collected on grapevine, for three out of the four scoring dates. Aggressiveness of other groups of isolates on strawberry were similar to each other. Isolates from strawberry did not exhibit greater growth of necrosis on this host compared to other hosts.

On hydrangea leaves, no pairwise differences between isolates collected from different hosts were observed, including for isolates collected on this host.

On bramble leaves, isolates collected from bramble did not exhibit the best performance, at any scoring time. Isolates from tomato were more aggressive on bramble than both grapevine isolates (range of mean diameter necrosis: 0.36-12.7 mm; significant at all scoring times) and bramble isolates (range of mean diameter necrosis: 0.2-11.80; significant for 3 out of 6 scoring times).

On tomato leaves, isolates collected from this host exhibited the highest mean aggressiveness, for the three latest scoring dates. Aggressiveness of isolates collected on bramble and grapevine was significantly reduced on tomato (range of mean diameter necrosis: −0.25 - −13.20 mm for bramble isolates, − 0.33 - −13.60 mm for grapevine isolates; statistically significant at all dates but the earliest), compared to isolates collected on tomato. Isolates from bramble and grapevine hosts performed similarly on tomato leaves.

On grapevine leaves, aggressiveness of isolates was highly variable, as indicated by the high proportion of boxplot outliers. No significant difference in aggressiveness was observed among isolates from different hosts on grapevine, but the biggest mean necrosis diameters were measured for isolates originating from grapevine at the two latest scoring times. In summary, the results of cross-pathogenicity tests support host specialization of *B. cinerea* to tomato, and to a lower extent, to grapevine; the differences in aggressiveness between *B. cinerea* populations associated with tomato and grapevine were also observed on bramble leaves.

#### On berries

We assessed differences in aggressiveness between clusters using pathogenicity tests on tomato berries after 16 days of inoculation and grapevine berries, respectively after 16 and 12 days. Necrosis percentages for each cluster are reported in **Error! Reference source not found**. and differences between pairs of hosts of origin are presented in Table 4. Our statistical model detected a significant effect of the host of origin. On tomato berries, isolates collected from this host exhibited the highest mean aggressiveness (46.6% of mean necrosis). Aggressiveness of isolates collected on bramble (28% of mean necrosis) was significantly reduced on tomato compared to isolates collected on tomato. No significant difference was detected with isolates collected on grapevine (38.1% of mean necrosis).

On grapevine berries, percentages of necrosis were lower than on tomato berries. Isolates collected from this host exhibited higher mean aggressiveness (23.7% of mean necrosis) than isolates collected on tomato (12.2%, *P*-value=0.064) but lower mean aggressiveness than isolates collected on bramble (33.3%, *P*-value=0.052).

## Discussion

### *Host specialization in* B. cinerea *populations infecting grapevine and tomato*

Several previous studies investigating the population structure of *B. cinerea* proposed that host plant should be considered as a potent factor structuring populations of the pathogen, based on observed associations between patterns of population subdivision and the host of origin of isolates (see review and examples in Walker, 2016). In a previous (Walker et al., 2015) and the present study, we confirmed that most of the genetic variation segregating in B. cinerea populations is explained by hosts, and that the effect of geography is weaker. Two of the clusters identified were associated with tomato and grapevine, respectively, while other clusters were observed on all hosts but tomato, although in different frequencies. Cross-pathogenicity experiments were consistent with analyses of population subdivision, with patterns of quantitative pathogenicity consistent with specialization of tomato isolates to tomato host, and specialization of grapevine isolates to grapevine hosts. This pattern of quantitative pathogenicity was observed after inoculation on leaves, but also in berry tests, which represents the type of symptoms more commonly found in the field.

A pattern of pathogenicity partially consistent with specialization was previously observed in indoor samples of *B. cinerea* collected on lettuce and tomato but differences in pathogenicity were not associated with population subdivision (Leyronas *et al*., 2015). To the best of our knowledge, this is the first report of host-associated ecological divergence in *B. cinerea*, with patterns of population structure mirroring patterns of quantitative pathogenicity. Indeed, the association between population structure and pathogenicity features suggests that host specialization contributes to reduce gene flow between populations associated with different hosts, due to the reduced viability of migrants and hybrids. The association between adaptation to hosts and barriers to gene flow would be favored by limited migration of gametes (especially non-mobile sclerotia developing on plant debris) between phases of selection by the host and sexual reproduction on (or inside) the host’s tissue.

Comparison of pairwise *Jost’s D* between clusters suggest that the tomato host induces greater population differentiation, *i.e*. greater reduction of gene flow between populations, compared to other hosts. This may rely on functional variations into the diverse and sophisticated biochemical arsenal of B. cinerea. Indeed, this pathogen shows a large repertoire of secreted proteins, secondary metabolites and small interfering RNA (siRNA) that act as effectors in the interactions with its hosts (Amselem et al., 2011; Mbengue et al., 2016; Reino et al., 2004; Siewers et al., 2005; Valero-Jiménez et al., 2019; Weiberg et al., 2013). The possible role of the different fungal effectors in host specialization remain to be investigated, as well as the possible role of the host phytoalexins. Indeed, contrasted pathogenic interactions between fungal pathogens (*e.g. Cladosporium fulvum* and *Fusarium oxysporum* f. sp. *lycopersici*) and tomato were explained by variation of tomatine content produced by the host and the production of tomatinase, an enzyme produced by the pathogen and able to detoxify the saponin toxin produced by tomato (Melton *et al*., 1998; Sarhan & Kiraly, 1981). Then, many molecular mechanisms could be involved in the incomplete host specialization of the grey mold fungus. Actually a recent genome-wide association mapping of *B. cinerea* showed that its virulence against a range of domesticated and wild tomatoes is highly polygenic and involves a diversity of mechanisms (Breeze, 2019; Soltis *et al*., 2019). Moreover, the previous observation that conidia of *B. cinerea* strains isolated from tomato have a higher germination rate on tomato cutin than conidia of strains isolated from grapevine (Cotoras & Silva, 2005) suggests that some of the molecular determinants underlying host specialization must be involved from the very early stage of the plant-pathogen interaction. Altogether, this pleads for elucidating the molecular determinants of host specialization in *B. cinerea* and to determine their specificity towards host plants, such as tomato but also within the Solanaceae family. From an evolutionary perspective, disentangling the molecular evolution of these determinants would contribute to unveil speciation in the *Botrytis* genus, in the light of the domestication of solanaceous plants. The biological material characterized in this study, as well as the high quality genomic resources available for *B. cinerea* should allow populations genomic analysis investigating the direction and density of selection exerted by the host, tomato in particular.

### *Lack of host specialization in* B. cinerea *populations infecting bramble, strawberry and hydrangea*

In this study we extended previous analyses to new hosts, with the aim to better characterize the pattern of host specialization in *B. cinerea*. We found no evidence for host specialization in isolates of *B. cinerea* collected on hydrangea in outdoor nurseries, neither from population subdivision nor from cross-pathogenicity tests. However, our capacity to detect ecological divergence in pathogens from hydrangea may have been limited by the small number of isolates collected on this host, associated with a relatively high proportion of clones. Further prospections may help determining the pathogenicity profile of hydrangea-associated isolates, in particular in environmental conditions favoring *Botrytis* population isolation in hydrangea cropping, such as in cold climatic chambers used to keep plants during vernalization.

The lack of population cluster specifically associated with strawberry was surprising, since Leroch et al. (2013) identified using multiple-gene sequencing, a novel clade of B. cinerea called group S, associated with strawberry in Germany. Using the detection marker designed by Leroch et al. (2013), we did detect a non-negligible proportion of group S isolates in our dataset (data not shown). However, group S isolates were not assigned into a specific cluster, suggesting that the pattern of population subdivision previously observed, and which led to the description of group S, may result from specific selection pressures such as resistance to fungicides, rather than from host specialization. Indeed, crops from our dataset, with the exception of bramble, were regularly sprayed with chemical fungicides, which might have favored the maintenance of group S strains in populations. The lack of specific interactions between strawberry and strawberry-collected isolates in the cross-pathogenicity test also supports this hypothesis.

The case of bramble-associated isolates is more puzzling, given that populations sampled from this host were differentiated from other populations in a previous study (Walker *et al*., 2015) and partially in this one, but tests did not reveal higher aggressiveness of isolates collected from bramble, compared to isolates from other hosts. Surprisingly, the difference in aggressiveness between tomato- and grapevine-collected strains was observed when inoculating on bramble, whereas it belongs to a botanical family, the Rosaceae, distant from Solanaceae and Vitaceae. However, plant susceptibility to *Botrytis cinerea* shows very little association to the evolutionary distances between the plant species (Caseys *et al*., 2018). Henceforth, theses aggressiveness variations could be linked to species-specific interactions. We may assume that the decrease in gene flow is not sufficient to induce host specialization on bramble, compared to tomato and grapevine. Considering host specialization as a dynamic process, adaptation to bramble might be only in the early stages, consistent with the recent domestication of blackberry and the restricted cultivation areas. In comparison, grapevine was domesticated more than 2000 years ago and tomato more than 400 years ago and both represent large cultivation areas worldwide.

### Concluding remarks

Our study shows that *B. cinerea* is a model of polyphagous fungal plant populations, whose populations are often comprised of generalist individuals, but can also be specialized to particular hosts (*e.g*. on tomato). While revealing key aspects of the process of adaptive divergence, understanding host specialization in *B. cinerea* may also help forecasting disease emergence and implementing sound management strategies. Indeed, host specialization suggests synchrony between host and pathogen development, determining time-shifted epidemics onsets in sympatric hosts within a landscape. Some cultivated hosts may also “filter” some specialized populations, within the collection of generalist and specialist individuals constituting *B. cinerea*. Limited gene flow between populations specialized on different crops may then be accentuated by optimizing spatial crop organization within an agricultural area. Limiting the density of putative high-risk host plants (with low host specialization) in the vicinity of high value crops may help decreasing disease propagation, as well as delaying fungicide resistance evolution.

## Experimental procedures

### Sample collection – Population genetics

*Botrytis cinerea* samples were collected in four areas of France (Alsace, Champagne, Loire Valley and Provence), with one to five collection sites per area (**Error! Reference source not found**.). Collection sites were 1 to 173 km apart within areas, and 180 to 703 km apart between areas. Populations from Champagne and Provence were chosen as a subset of a previously published dataset (Walker *et al*., 2015), keeping the sample size similar to the number of newly characterized strains (n=331) to avoid biases related to unbalanced sample size. Populations from Alsace and the Loire Valley (n=350) were collected specifically for the present study and include hosts that have not been previously characterized. Samples were collected in June in years 2002, 2005, 2006, 2007, 2014 and 2015. Spring is favorable to the sampling of B. cinerea populations from various hosts, as spring conditions facilitate primary infections by ascospores and winter-surviving macroconidia. Primary infections can occur on flowers and young ovaries by penetrating through scars left by floral pieces. Samples were collected from five different hosts: (i) strawberry (*Fragaria x ananassa*; diseased berries), (ii) grapevine (Vitis vinifera; asymptomatic flower caps), (iii) wild blackberry (*Rubus fruticosus;* asymptomatic berries and floral pieces) from bushes surrounding vineyards and fields, (iv) hydrangea (*Hydrangea macrophylla*; symptomatic and asymptomatic flower buds and stems) from outside nurseries and (v) tomato (*Solanum lycopersicum;* symptomatic fruits or petioles; Error! Reference source not found.). Isolates from asymptomatic hosts (only strawberry and tomato hosts systematically exhibited symptomatic berries at the collection time) were collected after incubating dead flowers and/or young fruit with supposed *B. cinerea* infection in a moist chamber at room temperature until sporulation. Single spores were isolated for each sample and cultured on malt-yeast-agar (MYA; 20 g.L^-1^ malt extract, 5 g.L^-1^ yeast extract, 15 g.L^-1^ agar) at 23°C, under continuous light until sporulation. Six hundred and eighty one single-spored isolates were produced and kept as conidial suspensions in glycerol 20% at - 80°C until use.

### Microsatellite genotyping

DNA was extracted after 7 days of cultivation on MYA (malt extract 20 g.L^-1^; yeast extract 5 g.L^-1^; agar 15 g.L^-1^) medium at 21 °C in the dark. DNA was extracted using the “DNeasy^®^ 96 Plant Kit” (QIAGEN^®^) following manufacturer protocol. Strains were genotyped at eight microsatellite markers (Bc1, Bc2, Bc3, Bc4, Bc5, Bc6, Bc7 and Bc10; Fournier et al., 2002) in multiplex polymerase chain reactions by Eurofins Scientific (Nantes, France). Markers with overlapping allelic size ranges bore distinct fluorochromes, as detailed in Supplementary Information 2. Allele size was scored automatically after binning analysis. Allele scoring for Bc4 enabled to remove from the dataset isolates of *Botrytis pseudocinerea*, the sister species living in sympatry with *B. cinerea*, based on their private allele at this locus (Fournier *et al*., 2002).

### Population subdivision

Hierarchical AMOVA in Arlequin v3.5 (Excoffier & Lischer, 2010) was used to estimate population differentiation within and between samples grouped according to their area or host plant of origin. Three different methods were used to investigate population subdivision. The model-based Bayesian clustering implemented in the Structure V2.3.4 software was used first (Pritchard *et al*., 2000). Structure was run with admixture and correlated allele frequencies options, with 500,000 Markov Chain Monte Carlo iterations, following 100,000 burn-in steps, and results were processed with Structure Harvester (Earl & vonHoldt, 2012). Second, the Discriminant Analysis of Principal Components DAPC was used as a multivariate analysis of population subdivision implemented in the Adegenet 2.1.1 (Jombart & Ahmed, 2011) package in R 3.4.3. Third, the population structure was inferred using the maximum likelihood method Snapclust implemented in the Adegenet 2.1.1 package in R 3.4.3. Genetic differentiation between the clusters identified with Structure was measured using Weir & Cockerham’s *F*_*st*_ in Genepop 4.7.0 (Rousset, 2008) to choose an appropriate number of clusters (K) and *Jost’s D* in GenAlEx 6.5 (Peakall & Smouse, 2012).

### Genetic diversity and mode of reproduction

All calculations were performed on clusters identified with Structure, considering only genotypes with a membership coefficient greater than or equal to 0.7 in one cluster. Poppr 2.8.2 (Kamvar *et al*., 2014; Kamvar *et al*., 2015) was used to calculate the number of unique genotypes G and the clonal fraction 1 – *G/N* with *N* the number of individuals in the population, and the 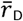 estimator of multilocus linkage disequilibrium. Allelic richness *Ar*, the mean number of alleles per locus, was estimated using a rarefaction method to standardize sample size in ADZE v1.0 (Szpiech *et al*., 2008). Gene diversity (He, called expected heterozygosity, in diploids) was estimated using Genetix (Belkhir *et al*., 1996-2004). Shannon’s diversity index, which captures the richness and evenness of allelic profiles, was computed using GenAlEx 6.5 (Peakall & Smouse, 2012).

### Sample collection – Pathogenicity tests

Cross-pathogenicity tests were carried out using single-spored isolates previously collected from sympatric grapevine, bramble and tomato hosts (Walker *et al.*, 2015), and single-spored isolates newly collected from strawberry, hydrangea and grapevine hosts (this study). Seventy-three isolates (7 to 33 isolates per host plant) were used in total (Supplementary Information 8). Only isolates with membership coefficient greater than or equal to 0.7 in the cluster they represent were used, and the laboratory reference strain B05.10 (Buttner *et al*., 1994), whose genome is sequenced, was used as a control. Strains were cultivated and kept as described in previous sampling procedure.

### Cross-pathogenicity tests

Isolates were cultivated on MYA at 23°C in the dark to prevent sporulation. Inoculations were performed using 1mm-diameter young mycelial plugs collected on the margin of the colonies, deposited on detached leaves (the mycelium facing the plant tissue), between leaf veins, and without wounding the leaf. Isolates were inoculated onto the following plants: tomato (cultivar Moneymaker; 2-3 week-old plantlets cultivated at 20°C with 8h dark), grapevine (cultivar Marselan; adult plants cultivated in greenhouse), bramble (wild plants collected in the surroundings of our laboratory), strawberry (cultivar Daroyal; one-year adult plants cultivated in greenhouse) and hydrangea (cultivar Leuchtfeuer; adult plants cultivated in greenhouse). Bean (*Phaseolus vulgaris*; cultivar Caruso; 2 week-old plantlets cultivated at 20°C with 8h dark) was used as a naive test host, differing from the previous collection hosts. Two sets of tests were conducted (Table 1): isolates from Champagne were inoculated onto grapevine, bramble, tomato and bean; isolates from Alsace and Pays de Loire were compared together on grapevine, strawberry and hydrangea.

**Table 1.** Hierarchical analysis of molecular variance (AMOVA) with host of collection or sampling location as grouping factor (upper tables) and host of collection nested within geographic origin as grouping factor (lower table). *P*-values in bold are significant at the 5% confidence level. d.f., degrees of freedom.

| Source of variation | d.f. | Sum of squares | Variance components | Percentage of variation | P-value | Fixation indices |
| --- | --- | --- | --- | --- | --- | --- |
| **Hosts** |  |  |  |  |  |  |
| Among hosts | 4 | 277 | 0.56 | 18.4 | **< 0.0001** | 0.184 |
| Within hosts | 696 | 1740 | 2.50 | 81.6 |  |  |
| Total | 700 | 2017 | 3.06 |  |  |  |
| **Geography** |  |  |  |  |  |  |
| Among areas | 5 | 104 | 0.16 | 5.5 | **< 0.0001** | 0.055 |
| Within areas | 695 | 1913 | 2.75 | 94.5 |  |  |
| Total | 700 | 2017 | 2.91 |  |  |  |
| **Host > Geography** |  |  |  |  |  |  |
| Among hosts | 4 | 277 | 0.49 | 16.2 | **< 0.0001** | 0.162 |
| Among hosts within areas | 11 | 112 | 0.19 | 6.1 | **< 0.0001** | 0.223 |
| Within populations | 685 | 1628 | 2.38 | 77.7 | **< 0.0001** | 0.073 |
| Total | 700 | 2017 | 3.06 |  |  |  |

**Table 2.**
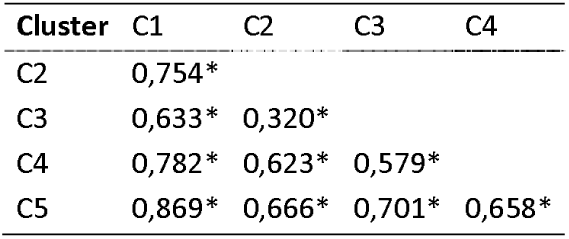
Pairwise *Jost’s D* between clusters inferred with STRUCTURE at *K*=5. Tomato, grapevine, grapevine, bramble and bramble were found as the main hosts for C1, C2, C3, C4 and C5, respectively (**Error! Reference source not found.**). Isolates were assigned into clusters when their membership coefficient was greater than 0.7 for a given cluster. Non-assigned isolates weren’t used in this analysis. *indicates that the pairwise *D* is significant at the 5% confidence levels.

| Cluster | C1 | C2 | C3 | C4 |
| --- | --- | --- | --- | --- |
| C2 | 0,754* |  |  |  |
| C3 | 0,633* | 0,320* |  |  |
| C4 | 0,782* | 0,623* | 0,579* |  |
| C5 | 0,869* | 0,666* | 0,701* | 0,658* |

**Table 3.** Results of cross-pathogenicity leaf tests observed over post-inoculation time for *B. cinerea* isolates representative of their host of collection. Isolates collected on strawberry, hydrangea, grapevine, bramble and tomato (hosts of origins, in columns) and a reference isolate used as a control were inoculated on strawberry, hydrangea, bean, bramble, tomato and/or grapevine leaves (test hosts, in row). Necrosis were scored between 2 and 16 days after inoculation, depending on the test host. ^a^ *P*-value indicating if lesion size according to host of origin was significantly different, on a given test host. ^b^ Significance of the *P*-value, indicating if test conditions (test, leaf and box repeat) significantly interacted with the lesion size on a given test host. *c* Values indicate the mean difference of necrosis diameter (in mm) between isolates collected on a host (2^nd^ row) compared to isolates collected on a second one (3^rd^ row), on given test host. Either positive or negative, these values indicate on which test host a given group of isolates had better growth. .,*, ** and *** indicates that the P-value is significant at the 10%, 5%, 1% and 1% confidence levels, respectively. - indicates that this interaction was not tested.

| Test host | Days after inoculation | Effect of the host of collection<sup>a</sup> | Test effect<sup>b</sup> | Control vs. |  |  |  |  | Strawberry vs. |  |  |  | Hydrangea vs. |  |  | Bramble vs. |  | Tomato |
| --- | --- | --- | --- | --- | --- | --- | --- | --- | --- | --- | --- | --- | --- | --- | --- | --- | --- | --- |
|  |  |  |  | Strawberry | Hydrangea | Bramble | Tomato | Grapevine | Hydrangea | Bramble | Tomato | Grapevine | Bramble | Tomato | Grapevine | Tomato | Grapevine | Grapevine |
| Leaf tests : Pairwise difference in mean necrosis diameter for individuals from various hosts of collection (mm)<sup>c</sup> |  |  |  |  |  |  |  |  |  |  |  |  |  |  |  |  |  |  |
| Strawberry | 2 | 0.005** | *** | 0.49 | 1.00 | - | - | 0.25 | 0.51 | - | - | -0.25 | - | - | -0.75** | - | - | - |
|  | 3 | 0.057. |  | 0.27 | 1.32 | - | - | -0.27 | 1.05 | - | - | -0.53 | - | - | -1.58* | - | - | - |
|  | 4 | 0.3 |  | -0.07 | 1.48 | - | - | -1.34 | 1.55 | - | - | -1.27 | - | - | -2.82 | - | - | - |
|  | 5 | 0.035* |  | -1.20 | 2.33 | - | - | -2.14 | 3.53 | - | - | -0.94 | - | - | -4.47* | - | - | - |
| Hydrangea | 2 | 0.46 | . | 0.43 | 0.36 | - | - | 0.44 | -0.07 | - | - | 0.017 | - | - | 0.08 | - | - | - |
|  | 3 | 0.31 | ** | 1.74 | 0.61 | - | - | 1.81 | -1.13 | - | - | 0.08 | - | - | 1.20 | - | - | - |
|  | 4 | 0.17 | *** | 2.82 | -0.22 | - | - | 2.77 | -3.04 | - | - | -0.05 | - | - | 2.99 | - | - | - |
|  | 5 | 0.17 | *** | 4.09 | -0.61 | - | - | 3.81 | -4.70 | - | - | -0.28 | - | - | 4.42 | - | - | - |
| Bean | 2 | 0.3 | *** | - | - | 2.41 | 1.65 | 1.46 | - | - | - | - | - | - | - | -0.75 | -0.95 | -0.19 |
|  | 3 | 0.3 | *** | - | - | 5.57 | 3.89 | 3.81 | - | - | - | - | - | - | - | -1.67 | -1.76 | -0.08 |
|  | 4 | 0.28 | *** | - | - | 8.30 | 4.86 | 5.93 | - | - | - | - | - | - | - | -3.44 | -2.36 | 1.08 |
|  | 5 | 0.099. |  | - | - | 5.93 | -1.00 | 7.54 | - | - | - | - | - | - | - | -6.93 | 1.60 | 8.54 |
| Bramble | 2 | 0.049* |  | - | - | 0.27 | 0.05 | 0.41 | - | - | - | - | - | - | - | -0.2 | 0.14 | 0.36* |
|  | 3 | 0.029* | *** | - | - | 0.76 | 0.11 | 0.79 | - | - | - | - | - | - | - | -0.65 | 0.03 | 0.68* |
|  | 4 | 0.0031** | *** | - | - | 2.58 | 0.29 | 2.39 | - | - | - | - | - | - | - | -2.29* | -0.20 | 2.10** |
|  | 5 | 0.0084** | *** | - | - | 3.84 | 0.57 | 3.78 | - | - | - | - | - | - | - | -3.28* | -0.06 | 3.21* |
|  | 8 | 0.005** |  | - | - | 9.04 | -2.78 | 9.96 | - | - | - | - | - | - | - | -11.80* | 0.92 | 12.70** |
|  | 9 | 0.044* | *** | - | - | 8.1 | 0.49 | 9.15 | - | - | - | - | - | - | - | -7.60 | 1.05 | 8.65* |
| Tomato | 2 | 0.091. | *** | - | - | 0.33 | 0.08 | 0.41 | - | - | - | - | - | - | - | -0.25 | 0.08 | 0.33. |
|  | 3 | 1.1e-06*** | *** | - | - | 5.54* | 1.24 | 5.40* | - | - | - | - | - | - | - | -4.31*** | -0.15 | 4.16*** |
|  | 4 | 0.00024*** | *** | - | - | 9.37 | 1.10 | 9.67 | - | - | - | - | - | - | - | -8.27** | 0.31 | 8.57*** |
|  | 5 | 0.0006*** | *** | - | - | 15.70 | 2.44 | 16.00 | - | - | - | - | - | - | - | -13.20** | 0.34 | 13.60*** |
| Grapevine | 2 | 0.4 | *** | 0.20 | 0.08 | 0.42 | 0.21 | 0.35 | -0.12 | 0.22 | 0.02 | 0.15 | 0.336 | 0.132 | 0.27 | -0.20 | -0.07 | 0.13 |
|  | 3 | 0.26 | *** | 1.33 | 1.11 | 1.37 | 0.65 | 0.67 | -0.22 | 0.04 | -0.68 | -0.66 | 0.26 | -0.456 | -0.43 | -0.72 | -0.69 | 0.02 |
|  | 4 | 0.12 | *** | 2.36 | 2.25 | 3.15 | 1.45 | 1.14 | -0.11 | 0.79 | -0.90 | -1.22 | 0.895 | -0.798 | -1.11 | -1.69 | -2.01 | -0.31 |
|  | 5 | 0.037* | *** | 3.82 | 4.09 | 4.03 | 1.93 | 0.73 | 0.27 | 0.22 | -1.89 | -3.09 | -0.0541 | -2.16 | -3.36 | -2.11 | -3.31 | -1.20 |
|  | 9 | 0.15 | *** | - | - | 14.2 | 8.74 | 4.76 | - | - | - | - | - | - | - | -5.50 | -9.47 | -3.98 |
| Berry tests : Pairwise log odds ratio from various hosts of collection |  |  |  |  |  |  |  |  |  |  |  |  |  |  |  |  |  |  |
| Tomato fruit | 16 | 0.02401 * | - | - | - | 0.52 | -0.77 | -0.24 | - | - | - | - | - | - | - | -1.30* | -0.77 | 0.52 |
| Grapevine | 12 | 0.000663 *** | *** | - | - | -0.57 | 0.74 | 0.097 | - | - | - | - | - | - | - | 1.31*** | 0.66 | -0.64 |

**Table 4.**
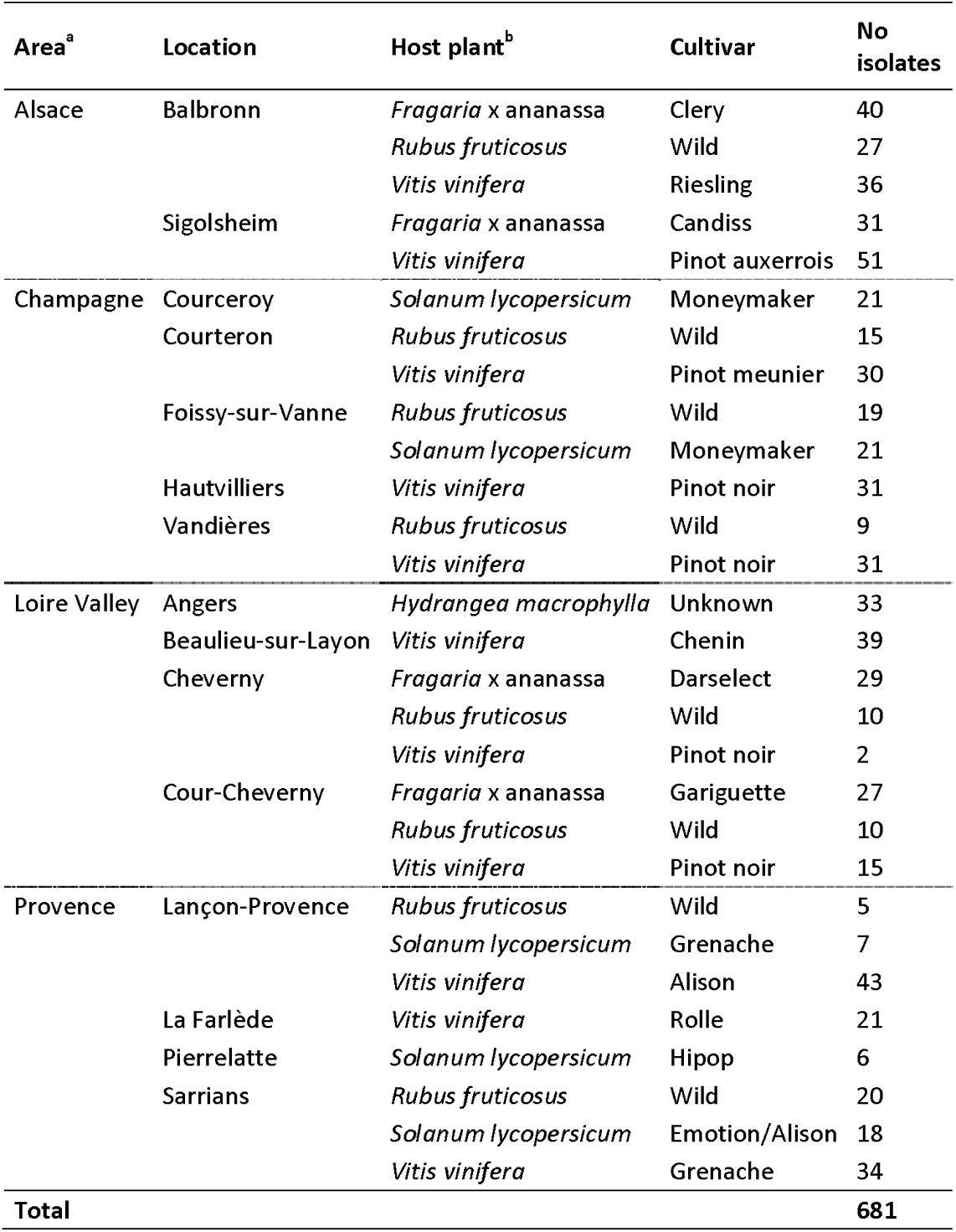
Populations of *B. cinerea* collected in four French areas, on four host plants between 2002 and 2015. ^a^ Isolates were sampled at multiple dates between 2002 and 2015 in Alsace (June 2015), Champagne and Provence (September 2005 to June 2007) and Loire Valley (June 2002 to June 2015). ^b^ Isolates collected on bramble, grapevine and hydrangea were retrieved from asymptomatic flower parts or young fruits. Isolates from tomato and strawberry were isolated from symptomatic fruits.

Inoculations were performed between May and July, when grapevine, strawberry and hydrangea have young leaves. Leaves were detached with sterilized scissors on the day of inoculation, and they were chosen to be as similar as possible in terms of size, age, and stalk position. Each plant species was challenged with all isolates within a week. Experiments were repeated twice on bean, bramble, strawberry and hydrangea, and four repeats were carried out on tomato and grapevine. For each test, ten plugs per isolate were randomly distributed across leaves; the reference isolate B05.10 was also deposited as a control on each leaf. The number of plugs (from different isolates) on each leaf differed across hosts, as they differ in leaf surface: 5 per tomato leaf, 3 per bean leaflet, 3 per grapevine and strawberry leaf, and 4 per hydrangea leaf. Inoculated leaves were incubated in 30 x 30 x 2 cm transparent plastic boxes, over paper moisturized with 40 ml sterile water and these moist chambers were incubated in climate chambers at 23°c with 8h of daily obscurity. Two diameters of each lesion surrounding plug inoculation were measured daily, during between 2 to 9 days depending on the plant species and the speed of necrosis maturation. Lesions were scored until they coalesced with neighboring lesions. For each test and each lesion, plug, leaf and box numbers were recorded in order to include this information in statistical tests. Altogether, the dataset on leaves was comprised of 25,863 measurements of lesion diameter.

For berry tests, isolates were cultivated on MYA at 23°C under continuous light to promote sporulation. Only isolates collected from tomato, grapevine and bramble were used in berry tests (a subset of isolates described in Supplementary Information 8). Ten-day old cultures were used to produce spore suspensions in sterile water at 5.10^5^ spores/ml, the day of the test. 10 µl droplets of spore suspension were deposited on the unwounded apical side of grapevine (white cultivar Mélissa) and tomato (unknown cultivar of cherry tomato) ripe berries. Berries were previously cleaned twice for 5 min in sterile water with 10% Tween80, rinsed two times for 5 min in pure water and dried before use. Ten inoculated berries per isolate were stabilized using individual gaskets in a 24 x 18 x 10 cm transparent plastic box over paper moisturized with 20 ml sterile water. A set of ten non-inoculated berries was added in each test to detect natural contamination of the berries. These moist chambers were incubated as for leaf tests. The test was repeated two times for each isolate. The percentage of necrotic area was scored for each berry after 16 and 12 days for tomato and grapevine, respectively. Altogether, the dataset on berries was comprised of 1600 measurements of necrosis surface.

### Statistical analysis

Hosts of origin were tested for their effect on leaf necrosis. For each tested host and for each day after inoculation, a linear mixed model was used, with diameter of necrosis measured on leaf as the response variable, the host of origin as fixed effect and the isolate, leaf and test box as random effects (using the R packages lmer and lmerTest; Bates *et al*., 2015; Kuznetsova *et al*., 2017). The mean of diameter of necrosis was compared between hosts of origin of isolates using a *post-hoc* Tukey test (using the R package emmeans; Lenth *et al*., 2019).

The effect of hosts of origin was further tested on percentage of necrosis on fruit. A generalized linear mixed model with percentage of necrosis as the response variable, the host of origin and the test as fixed effects and isolate as a random effect was used for each tested host and for each day after inoculation. The necrosis percentage was compared between hosts of origin of isolates using a *post-hoc* Tukey test.

## Acknowledgements

AM was supported by a grant from the Doctoral School " Sciences du Végétal ", Université Paris-Saclay. We thank Marguerite Cuel, for preliminary data analysis of the cross-pathogenicity tests. We thank Claire Amiraux (Planète Légumes), Marie-Laure Panon (Comité Champagne), Jean-Marie Guichardon and Michel Badier (Chambre d’agriculture du Loire et Cher) and Elisabeth Fournier (UMR BGPI, INRA) for help with the sampling of *B. cinerea* isolates. We are also grateful to Thomas Perron (IRHS, INRA, Angers), the Vegepolys Pôle de Compétitivité, the Region Pays de la Loire (*Physi’Ho* project) and the local horticultors for providing the isolates from hydrangea.

---

**Supplementary Information 1.**
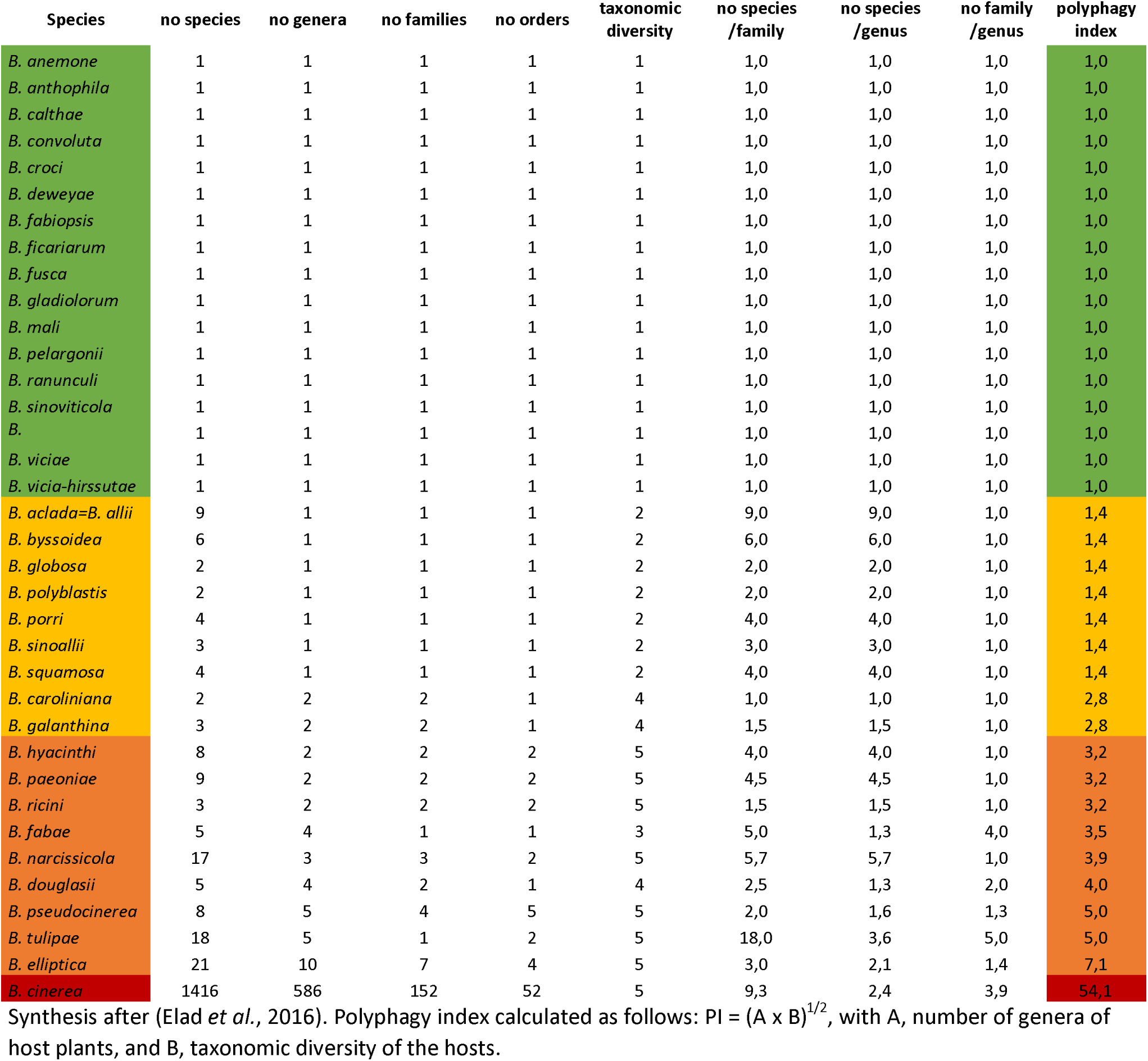
Polyphagy status of various *Botrytis* species.

**Supplementary Information 2.**
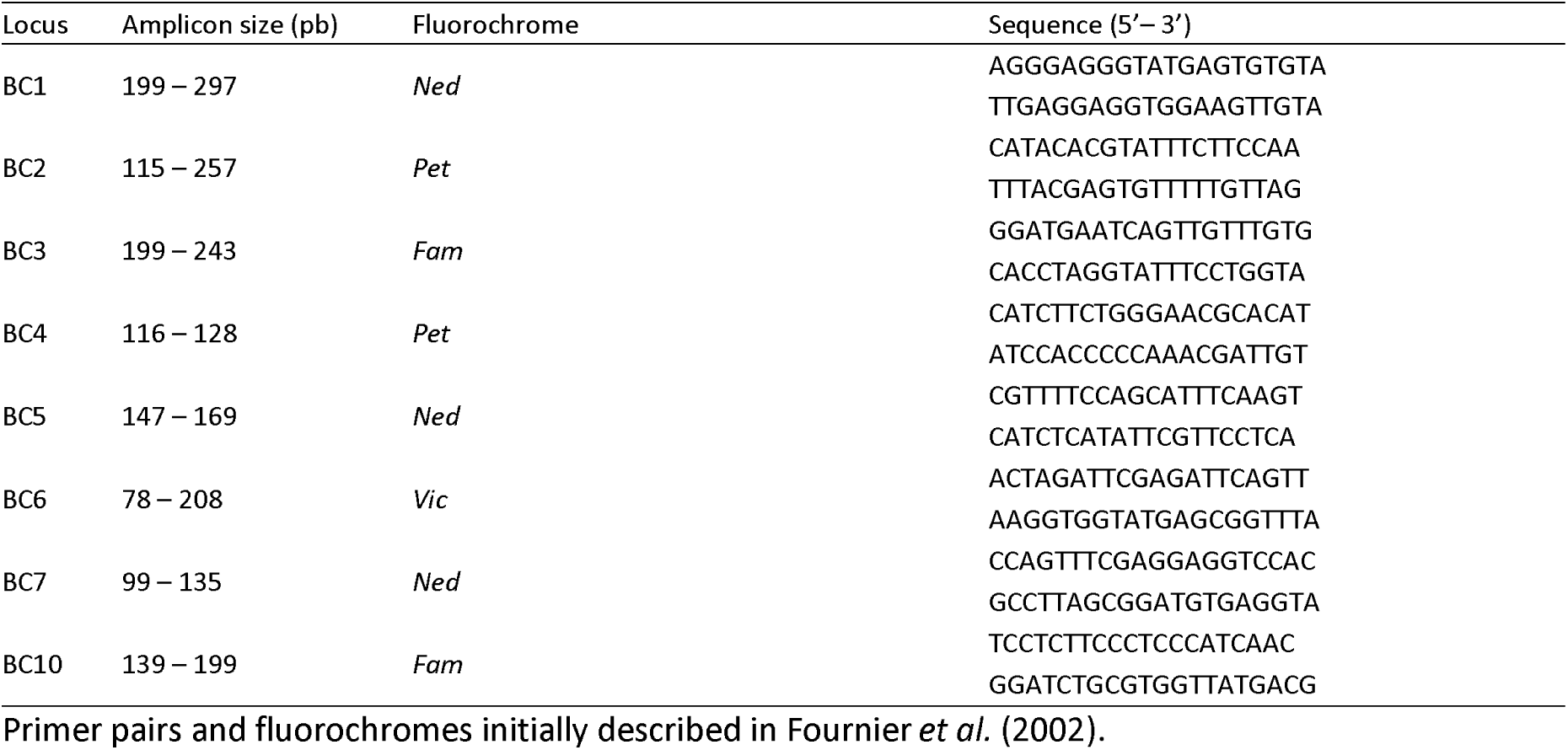
Primers, fluorochromes and PCR conditions for microsatellite genotyping.

5 µl of DNA (5 ng/µl) and 1.25 µl of each primer (2 µM) was used per multiplex PCR reaction, with the Taq polymerase Type-it (Qiagen), according to the manufacturer’s recommendations. The PCR program was:

- Initial denaturation: 94°C for 5 mn
- Elongation (35 cycles): 94°C 30s, 55°C 30s, 72°C 2mn
- Final elongation: 72°C 2mn

Amplicons were observed with the GeneScan and Genotyper programs (Life Technologies) after capillary electrophoresis achieved on an ABI3130xl sequencer, using the Liz500 size standard (Life Technologies).

**Supplementary Information 3.**
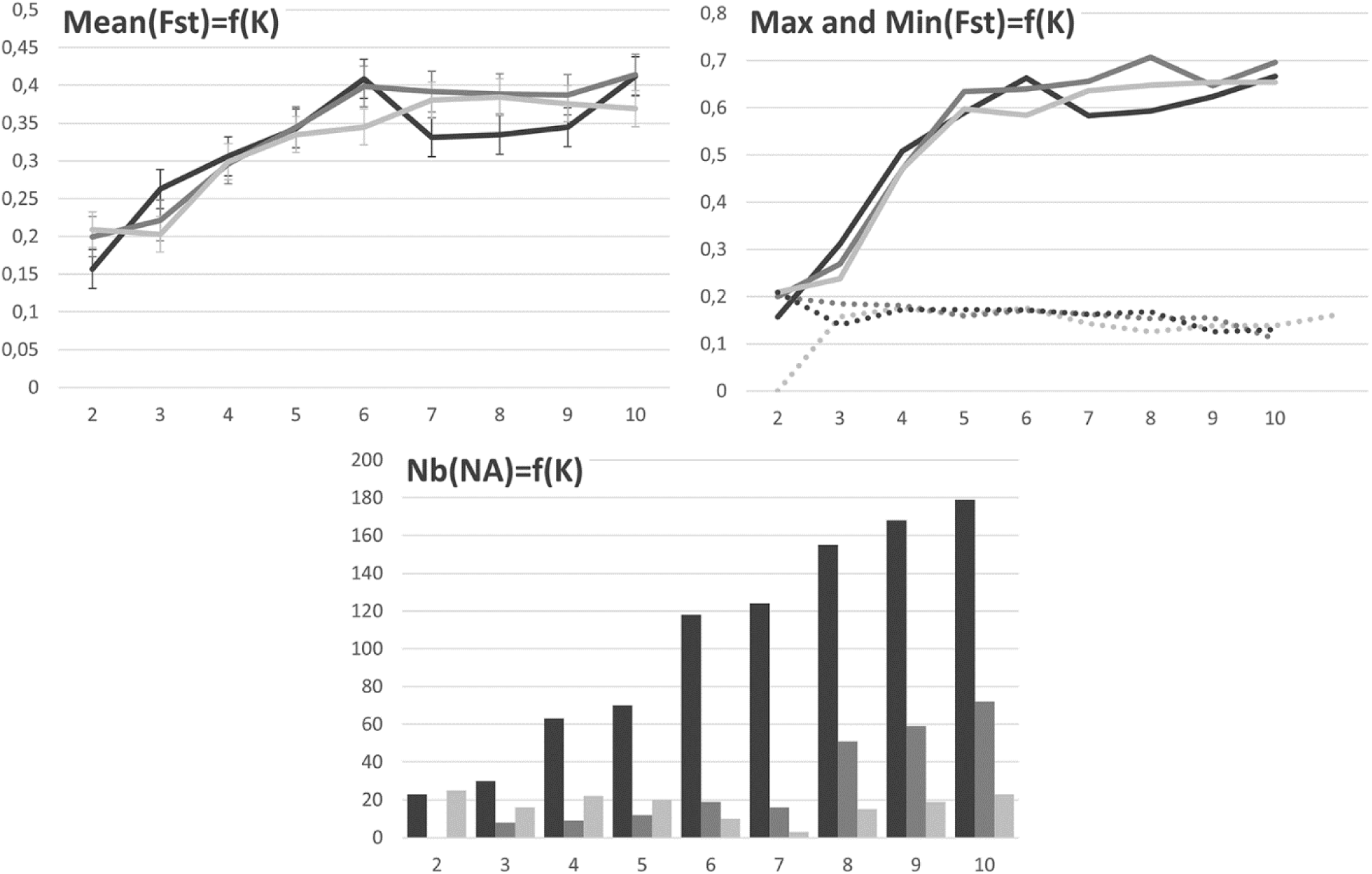
*F*_*st*_ and number of unassigned individuals with increasing *K* values.

Differentiation between clusters (*F*_*st*_) and number of ‘unassigned’ genotypes (‘assigned’ genotypes with >90% membership proportion or membership probability in a single group) as a function of the number of clusters *K* with Structure (black), DAPC (dark grey) and Snapclust (light grey) methods. The first graphic (upper left) represents the mean *F*_*st*_ between clusters along *K* increments. Vertical bars are the standard deviation. The second graphic (upper right) represents the maximum (solid line) and minimum (dotted line) values of *F*_*st*_ between clusters along increments.

**Supplementary Information 4.**
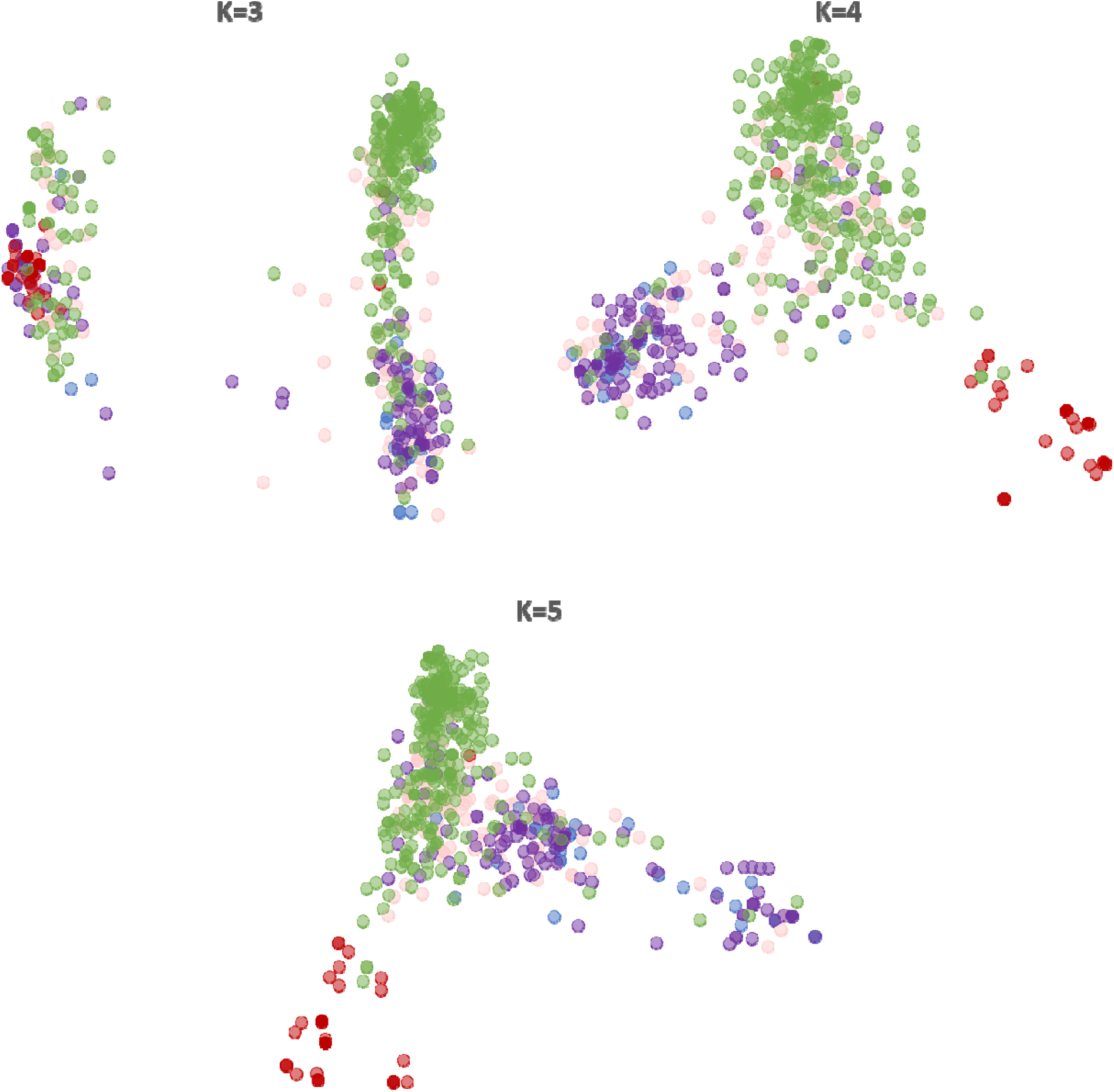
Scatterplots of DAPC assignations.

Each scatterplot represents the axes 1-2 of a DAPC assignation for values of K from 3 to 5. Individuals are colored according to their host of collection (*Solanum* in red, *Vitis* in green, *Rubus* in purple, *Hydrangea* in blue and *Fragaria* in pink).

**Supplementary Information 5.**
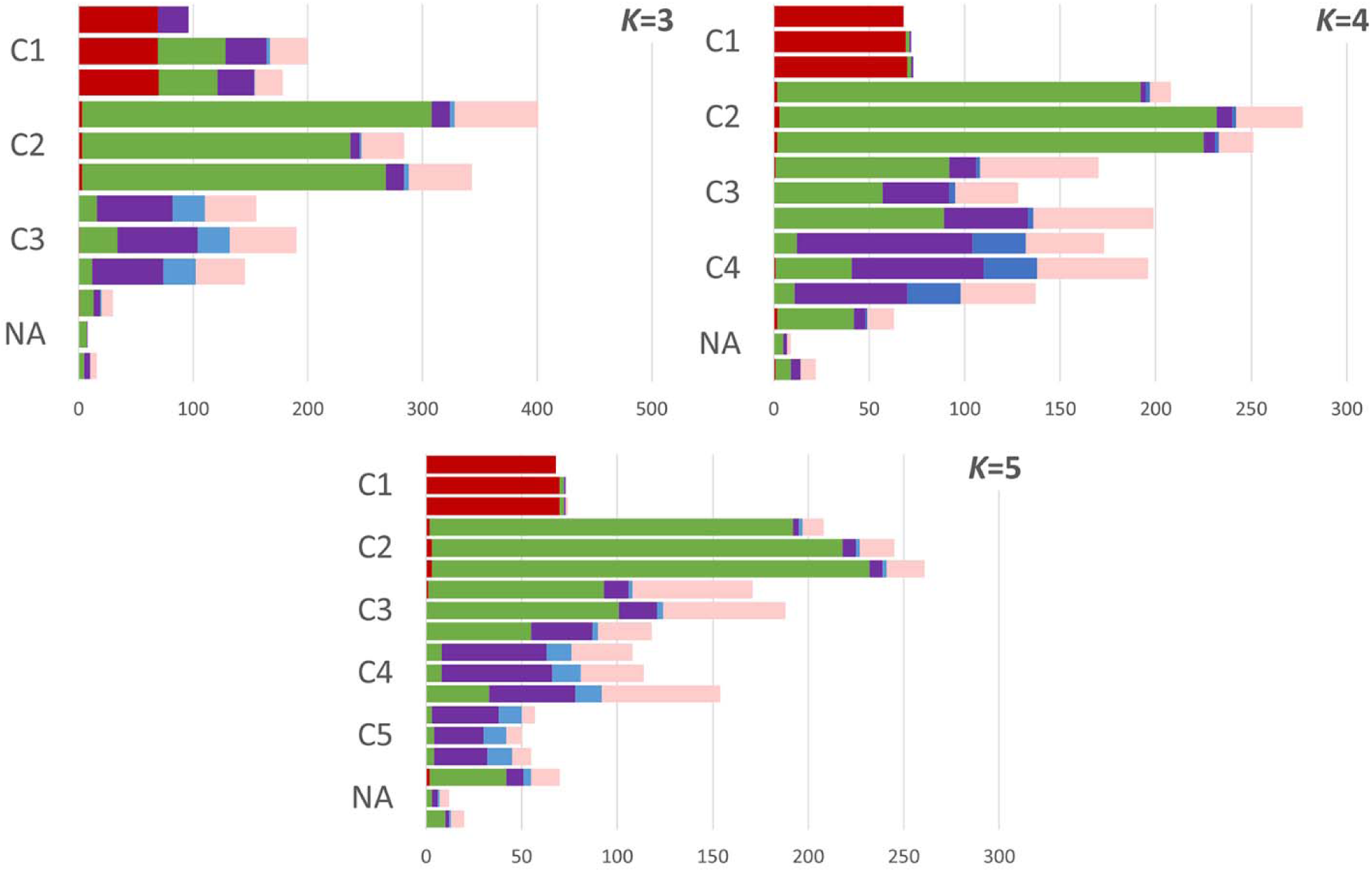
Host of origin of assigned individuals.

Each plot represent the proportion of individuals sampled from different hosts colored accordingly (*Solanum* in red, *Vitis* in green, *Rubus* in purple, *Hydrangea* in blue and *Fragaria* in pink) at *K* values from 3 to 5. The upper bar corresponds to the Structure assignation, the middle one to DAPC and lower one to Snapclust.

**Supplementary Information 6.**
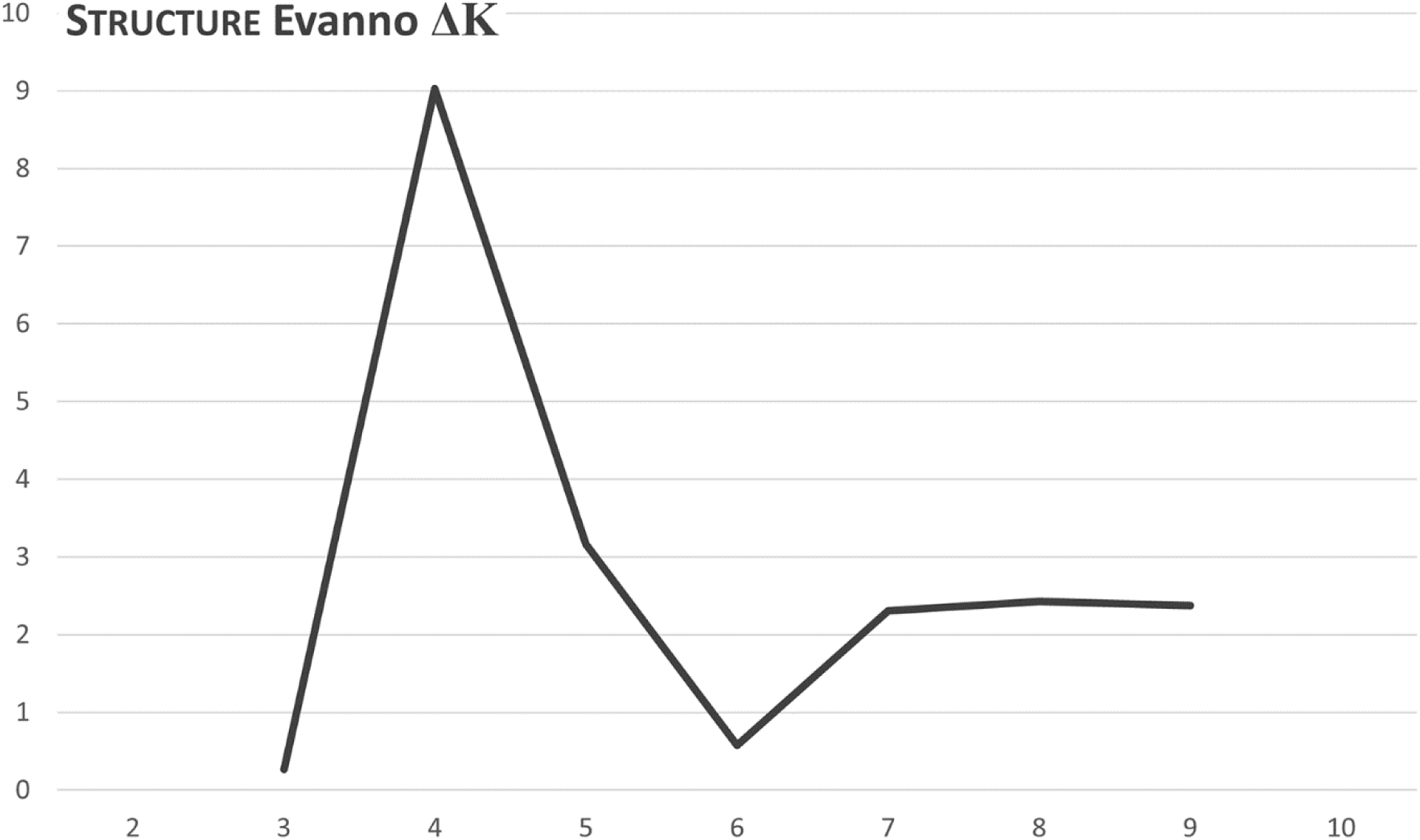
Evanno’s ΔK statistic.

The ΔK statistic (Evanno *et al*., 2005) is calculated on Structure results along *K* values from 2 to 10.

**Supplementary Information 7.**
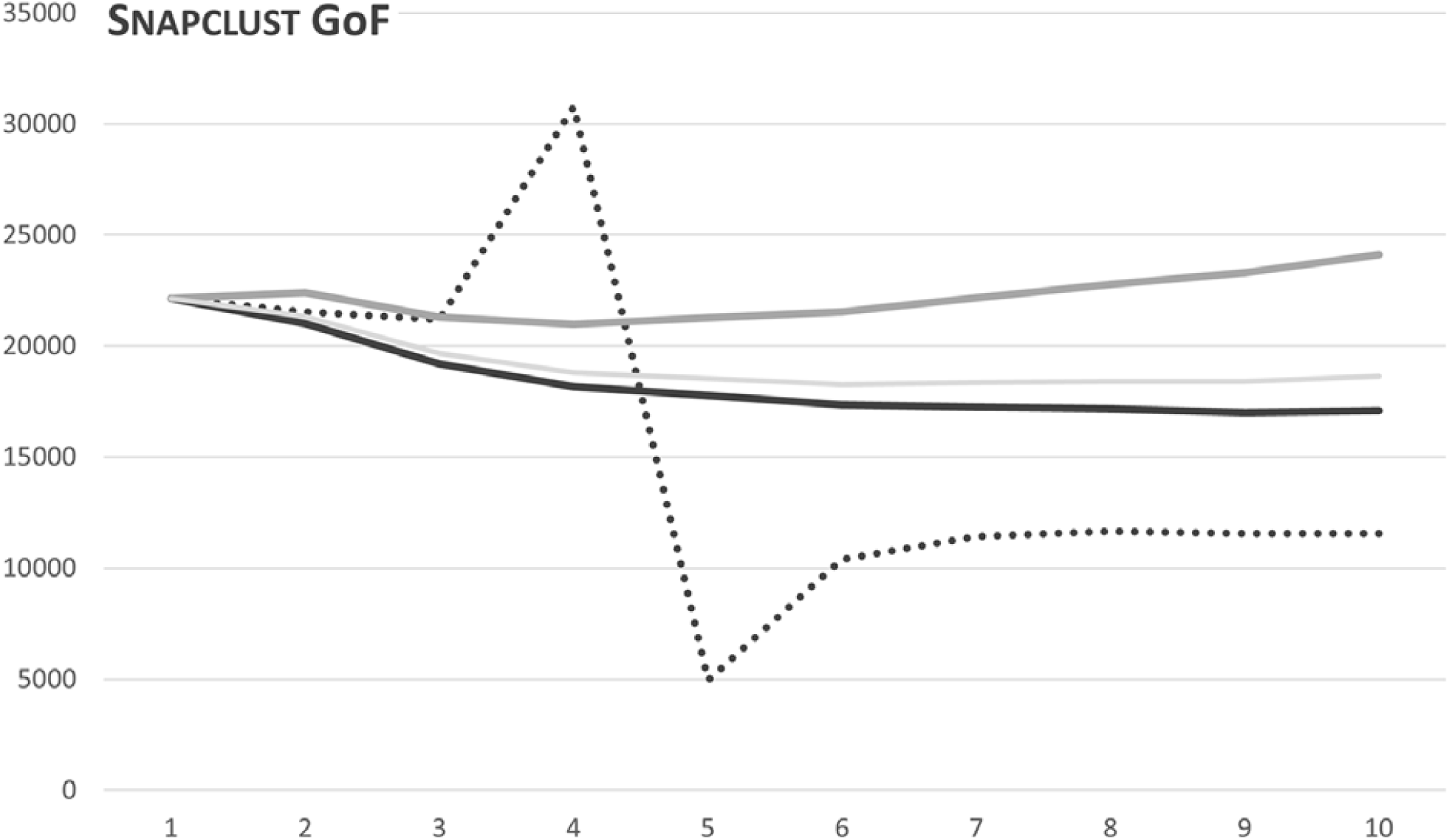
Goodness of fit statistics.

Each line corresponds to one of the goodness of fit statistic available with Snapclust (Jombart, 2008). The solid black line corresponds the AIC, the dotted black line to the AICc, the solid dark gray line to the BIC and the solid light gray line to the KIC.

**Supplementary Information 8.**
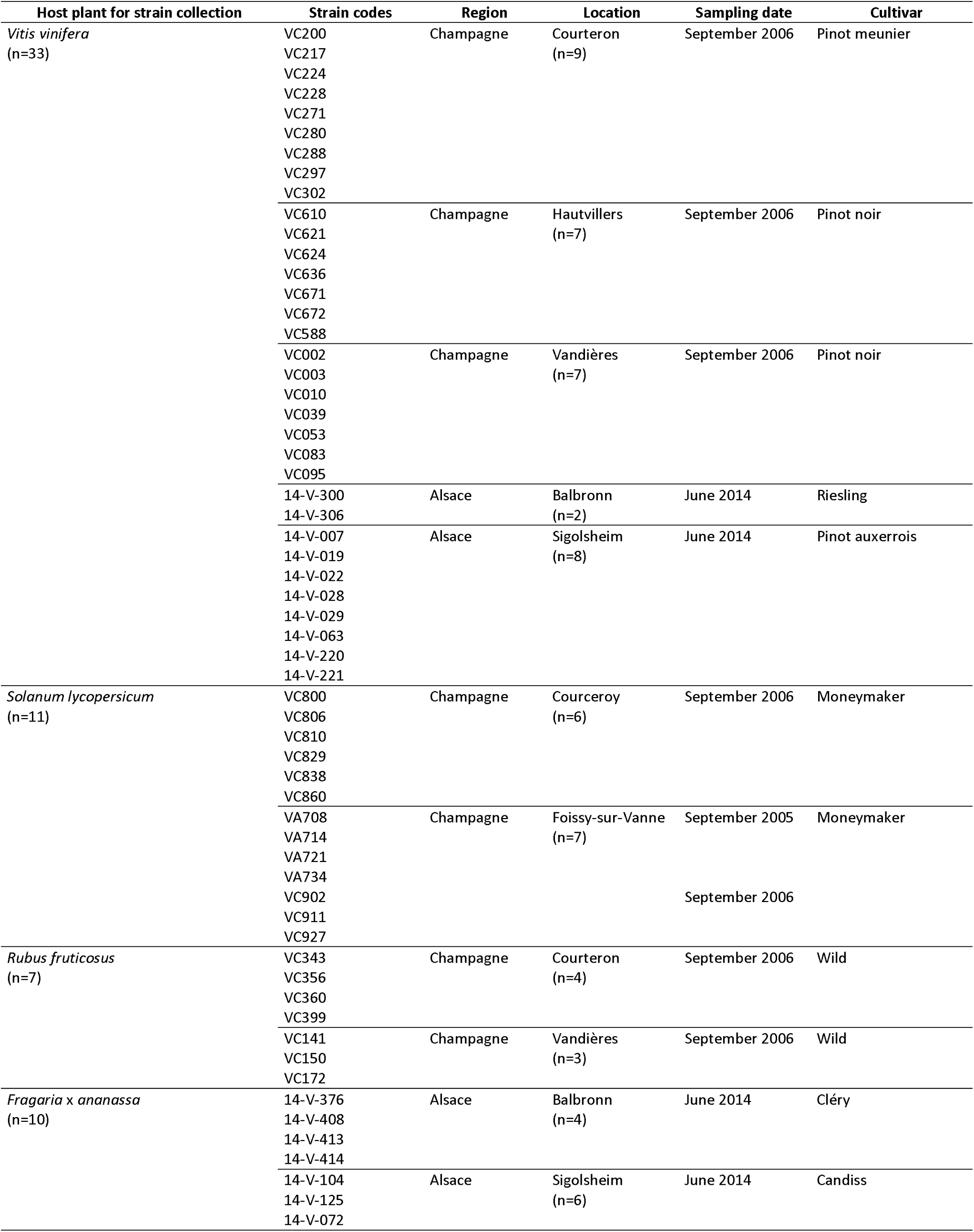
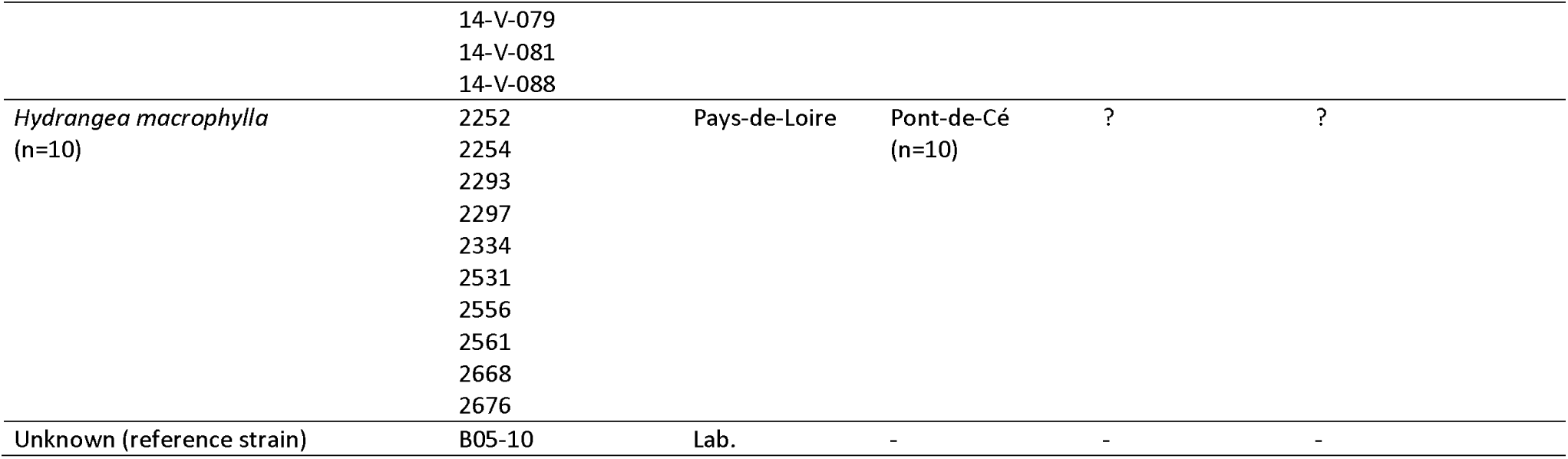
*Botrytis cinerea* strains and origin used to test host preference in microbiological tests.

