## Supplementary Information 1-8 for "The polyphagous plant pathogenic fungus *Botrytis cinerea* encompasses host-specialized and generalist populations"

### **Supplementary Information 1.** Polyphagy status of various *Botrytis* species.

| **Species** | **no species** | **no genera** | **no families** | **no orders** | **taxonomic diversity** | **no species /family** | **no species /genus** | **no family /genus** | **polyphagy index** |
| --- | --- | --- | --- | --- | --- | --- | --- | --- | --- |
| *B. anemone* | 1 | 1 | 1 | 1 | 1 | 1,0 | 1,0 | 1,0 | 1,0 |
| *B. anthophila* | 1 | 1 | 1 | 1 | 1 | 1,0 | 1,0 | 1,0 | 1,0 |
| *B. calthae* | 1 | 1 | 1 | 1 | 1 | 1,0 | 1,0 | 1,0 | 1,0 |
| *B. convoluta* | 1 | 1 | 1 | 1 | 1 | 1,0 | 1,0 | 1,0 | 1,0 |
| *B. croci* | 1 | 1 | 1 | 1 | 1 | 1,0 | 1,0 | 1,0 | 1,0 |
| *B. deweyae* | 1 | 1 | 1 | 1 | 1 | 1,0 | 1,0 | 1,0 | 1,0 |
| *B. fabiopsis* | 1 | 1 | 1 | 1 | 1 | 1,0 | 1,0 | 1,0 | 1,0 |
| *B. ficariarum* | 1 | 1 | 1 | 1 | 1 | 1,0 | 1,0 | 1,0 | 1,0 |
| *B. fusca* | 1 | 1 | 1 | 1 | 1 | 1,0 | 1,0 | 1,0 | 1,0 |
| *B. gladiolorum* | 1 | 1 | 1 | 1 | 1 | 1,0 | 1,0 | 1,0 | 1,0 |
| *B. mali* | 1 | 1 | 1 | 1 | 1 | 1,0 | 1,0 | 1,0 | 1,0 |
| *B. pelargonii* | 1 | 1 | 1 | 1 | 1 | 1,0 | 1,0 | 1,0 | 1,0 |
| *B. ranunculi* | 1 | 1 | 1 | 1 | 1 | 1,0 | 1,0 | 1,0 | 1,0 |
| *B. sinoviticola* | 1 | 1 | 1 | 1 | 1 | 1,0 | 1,0 | 1,0 | 1,0 |
| *B. sphaerosperma* | 1 | 1 | 1 | 1 | 1 | 1,0 | 1,0 | 1,0 | 1,0 |
| *B. viciae* | 1 | 1 | 1 | 1 | 1 | 1,0 | 1,0 | 1,0 | 1,0 |
| *B. vicia-hirssutae* | 1 | 1 | 1 | 1 | 1 | 1,0 | 1,0 | 1,0 | 1,0 |
| *B. aclada=B. allii* | 9 | 1 | 1 | 1 | 2 | 9,0 | 9,0 | 1,0 | 1,4 |
| *B. byssoidea* | 6 | 1 | 1 | 1 | 2 | 6,0 | 6,0 | 1,0 | 1,4 |
| *B. globosa* | 2 | 1 | 1 | 1 | 2 | 2,0 | 2,0 | 1,0 | 1,4 |
| *B. polyblastis* | 2 | 1 | 1 | 1 | 2 | 2,0 | 2,0 | 1,0 | 1,4 |
| *B. porri* | 4 | 1 | 1 | 1 | 2 | 4,0 | 4,0 | 1,0 | 1,4 |
| *B. sinoallii* | 3 | 1 | 1 | 1 | 2 | 3,0 | 3,0 | 1,0 | 1,4 |
| *B. squamosa* | 4 | 1 | 1 | 1 | 2 | 4,0 | 4,0 | 1,0 | 1,4 |
| *B. caroliniana* | 2 | 2 | 2 | 1 | 4 | 1,0 | 1,0 | 1,0 | 2,8 |
| *B. galanthina* | 3 | 2 | 2 | 1 | 4 | 1,5 | 1,5 | 1,0 | 2,8 |
| *B. hyacinthi* | 8 | 2 | 2 | 2 | 5 | 4,0 | 4,0 | 1,0 | 3,2 |
| *B. paeoniae* | 9 | 2 | 2 | 2 | 5 | 4,5 | 4,5 | 1,0 | 3,2 |
| *B. ricini* | 3 | 2 | 2 | 2 | 5 | 1,5 | 1,5 | 1,0 | 3,2 |
| *B. fabae* | 5 | 4 | 1 | 1 | 3 | 5,0 | 1,3 | 4,0 | 3,5 |
| *B. narcissicola* | 17 | 3 | 3 | 2 | 5 | 5,7 | 5,7 | 1,0 | 3,9 |
| *B. douglasii* | 5 | 4 | 2 | 1 | 4 | 2,5 | 1,3 | 2,0 | 4,0 |
| *B. pseudocinerea* | 8 | 5 | 4 | 5 | 5 | 2,0 | 1,6 | 1,3 | 5,0 |
| *B. tulipae* | 18 | 5 | 1 | 2 | 5 | 18,0 | 3,6 | 5,0 | 5,0 |
| *B. elliptica* | 21 | 10 | 7 | 4 | 5 | 3,0 | 2,1 | 1,4 | 7,1 |
| *B. cinerea* | 1416 | 586 | 152 | 52 | 5 | 9,3 | 2,4 | 3,9 | 54,1 |

Synthesis after (Elad *et al.*, 2016). Polyphagy index calculated as follows: PI = (A x B)^1/2^, with A, number of genera of host plants, and B, taxonomic diversity of the hosts.

### **Supplementary Information 2.** Primers, fluorochromes and PCR conditions for microsatellite genotyping.

| Locus | Amplicon size (pb) | Fluorochrome | Sequence (5’– 3’) |
| --- | --- | --- | --- |
| BC1 | 199 – 297 | *Ned* | AGGGAGGGTATGAGTGTGTA |
|  |  |  | TTGAGGAGGTGGAAGTTGTA |
| BC2 | 115 – 257 | *Pet* | CATACACGTATTTCTTCCAA |
|  |  |  | TTTACGAGTGTTTTTGTTAG |
| BC3 | 199 – 243 | *Fam* | GGATGAATCAGTTGTTTGTG |
|  |  |  | CACCTAGGTATTTCCTGGTA |
| BC4 | 116 – 128 | *Pet* | CATCTTCTGGGAACGCACAT |
|  |  |  | ATCCACCCCCAAACGATTGT |
| BC5 | 147 – 169 | *Ned* | CGTTTTCCAGCATTTCAAGT |
|  |  |  | CATCTCATATTCGTTCCTCA |
| BC6 | 78 – 208 | *Vic* | ACTAGATTCGAGATTCAGTT |
|  |  |  | AAGGTGGTATGAGCGGTTTA |
| BC7 | 99 – 135 | *Ned* | CCAGTTTCGAGGAGGTCCAC |
|  |  |  | GCCTTAGCGGATGTGAGGTA |
| BC10 | 139 – 199 | *Fam* | TCCTCTTCCCTCCCATCAAC |
|  |  |  | GGATCTGCGTGGTTATGACG |

Primer pairs and fluorochromes initially described in Fournier *et al.* (2002).

5 µl of DNA (5 ng/µl) and 1.25 µl of each primer (2 µM) was used per multiplex PCR reaction, with the Taq polymerase Type-it (Qiagen), according to the manufacturer’s recommendations. The PCR program was:

- Initial denaturation: 94°C for 5 mn
- Elongation (35 cycles): 94°C 30s, 55°C 30s, 72°C 2mn
- Final elongation: 72°C 2mn

Amplicons were observed with the GeneScan and Genotyper programs (Life Technologies) after capillary electrophoresis achieved on an ABI3130xl sequencer, using the Liz500 size standard (Life Technologies).

### Supplementary Information 3. *F_st_* and number of unassigned individuals with increasing *K* values.

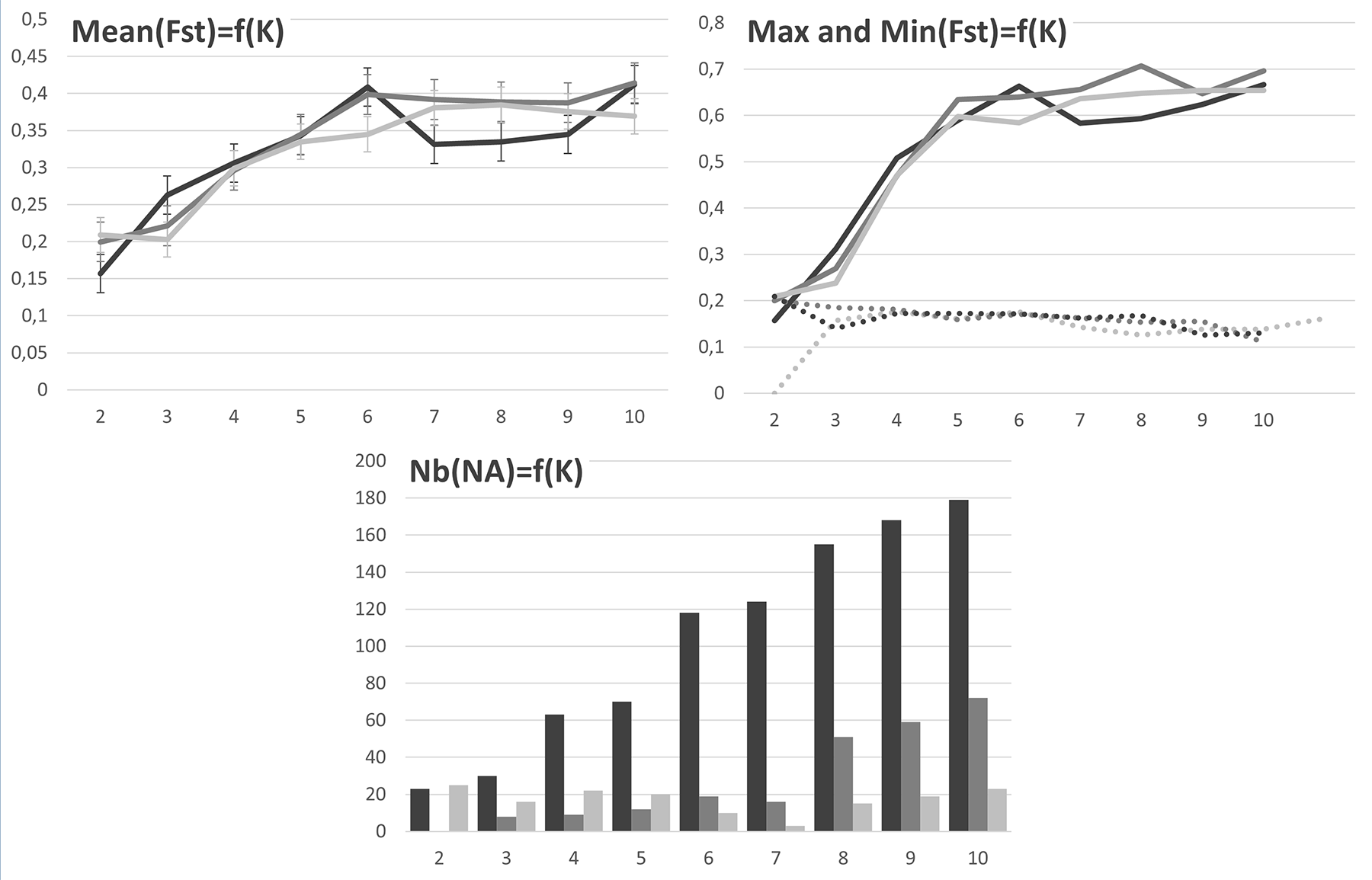

Differentiation between clusters (*F_st_*) and number of ‘unassigned’ genotypes (‘assigned’ genotypes with >90% membership proportion or membership probability in a single group) as a function of the number of clusters *K* with Structure (black), DAPC (dark grey) and Snapclust (light grey) methods. The first graphic (upper left) represents the mean *F_st_* between clusters along *K* increments. Vertical bars are the standard deviation. The second graphic (upper right) represents the maximum (solid line) and minimum (dotted line) values of *F_st_* between clusters along increments.

### Supplementary Information 4. Scatterplots of DAPC assignations.

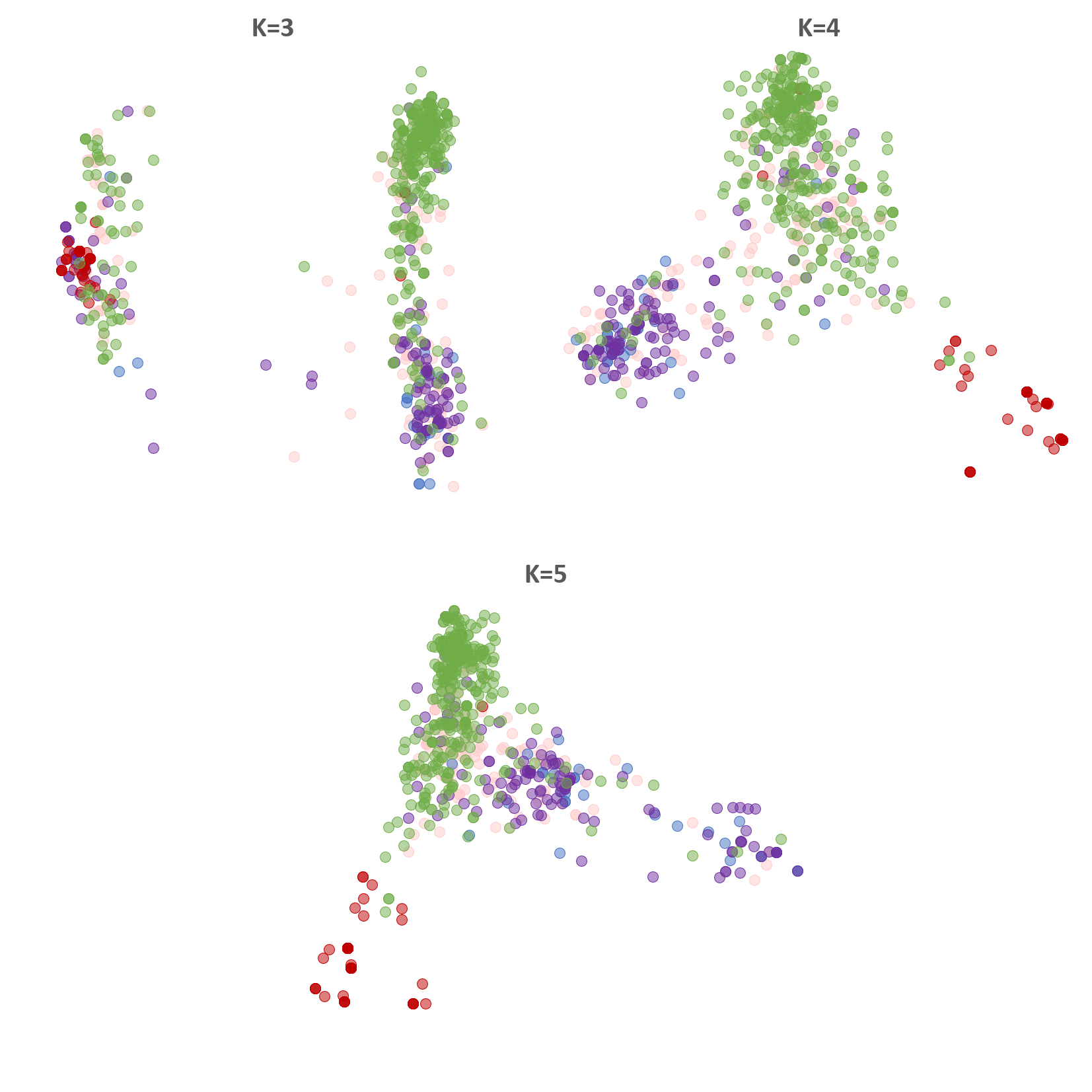

Each scatterplot represents the axes 1-2 of a DAPC assignation for values of *K* from 3 to 5. Individuals are colored according to their host of collection (*Solanum* in red, *Vitis* in green, *Rubus* in purple, *Hydrangea* in blue and *Fragaria* in pink**).**

### Supplementary Information 5. Host of origin of assigned individuals.

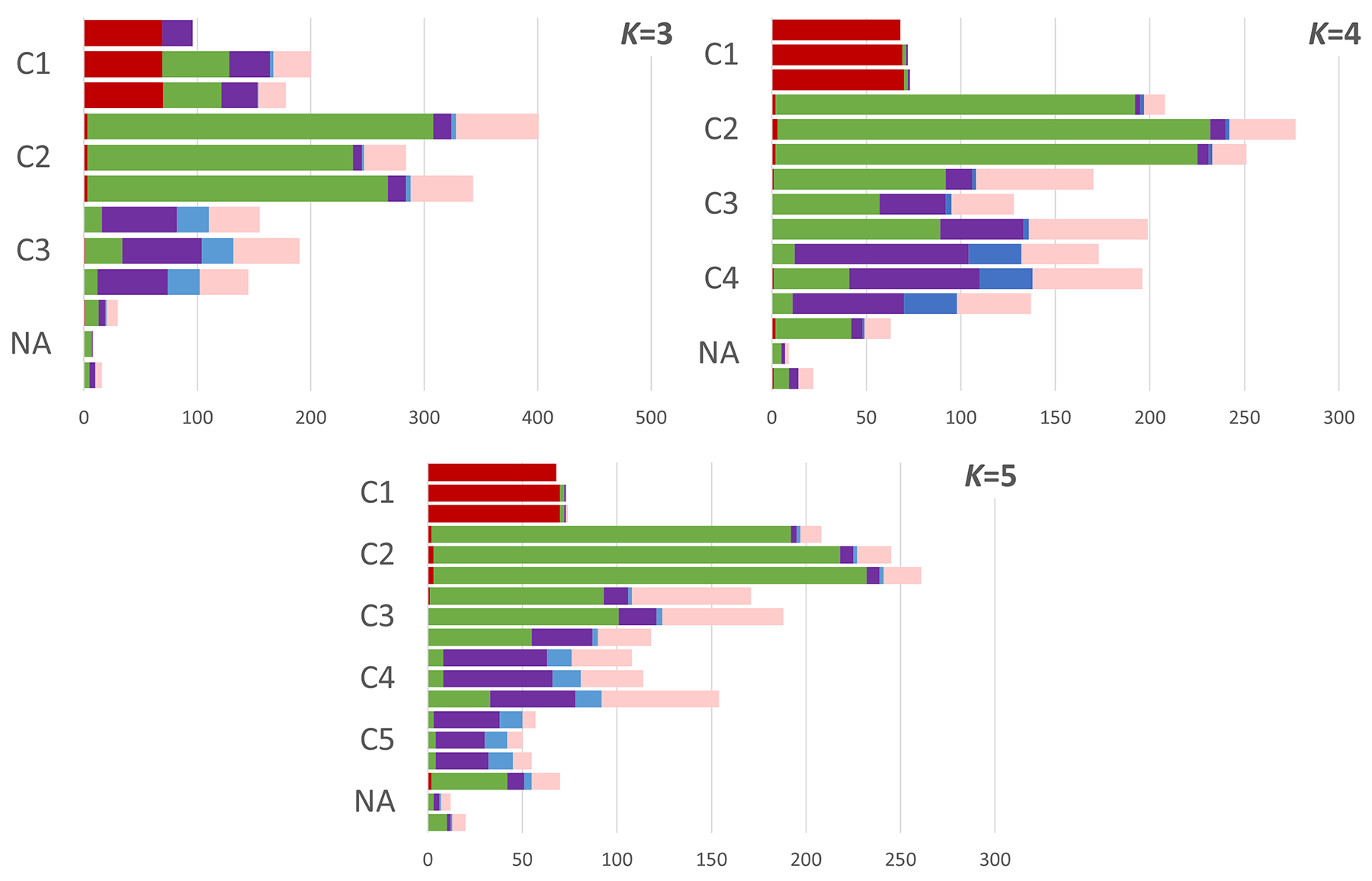

Each plot represent the proportion of individuals sampled from different hosts colored accordingly (*Solanum* in red, *Vitis* in green, *Rubus* in purple, *Hydrangea* in blue and *Fragaria* in pink) at *K* values from 3 to 5. The upper bar corresponds to the Structure assignation, the middle one to DAPC and lower one to Snapclust.

**Supplementary Information 6.** Evanno’s ΔK statistic.

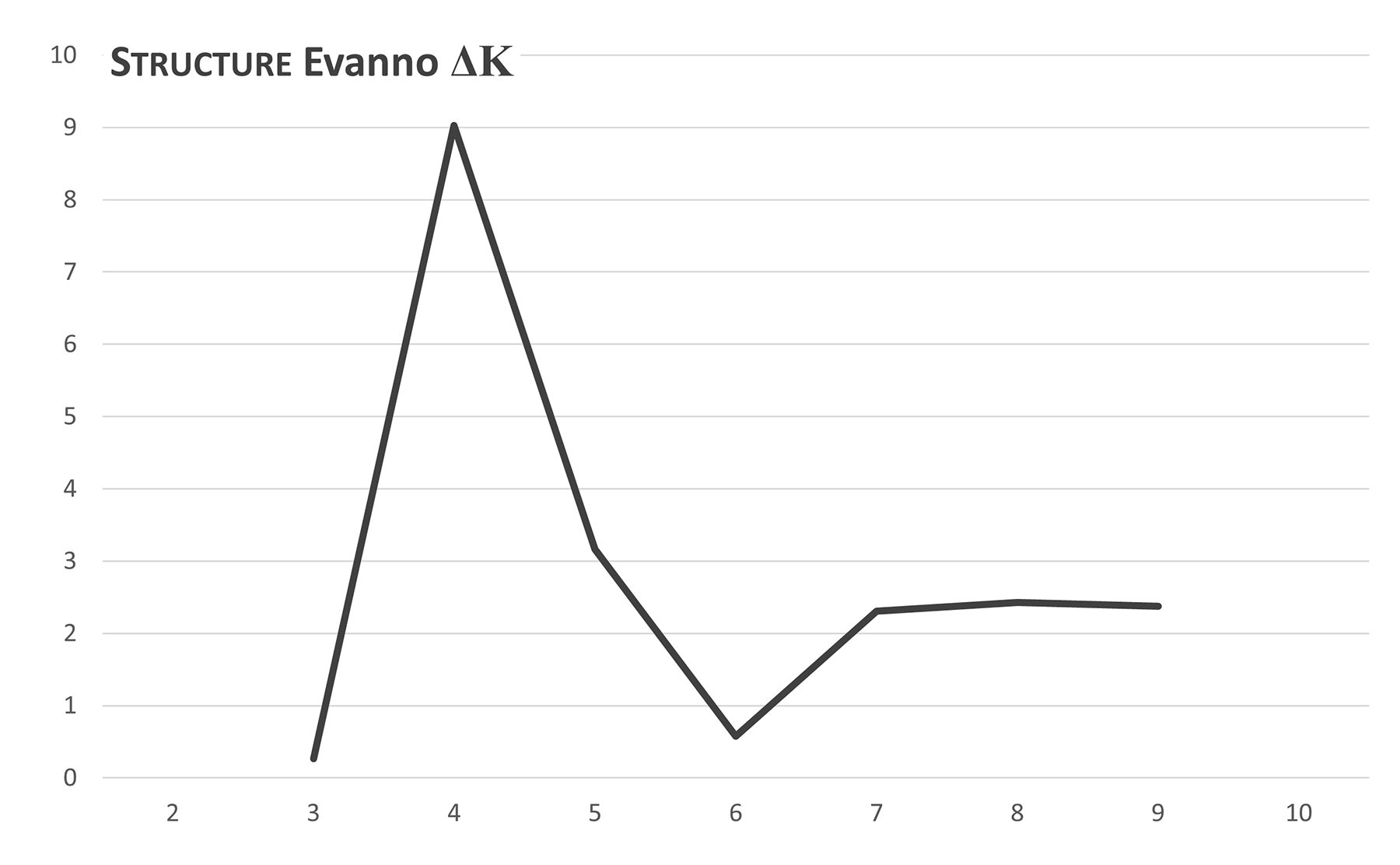

The ΔK statistic (Evanno *et al.*, 2005) is calculated on Structure results along *K* values from 2 to 10.

**Supplementary Information 7.** Goodness of fit statistics.

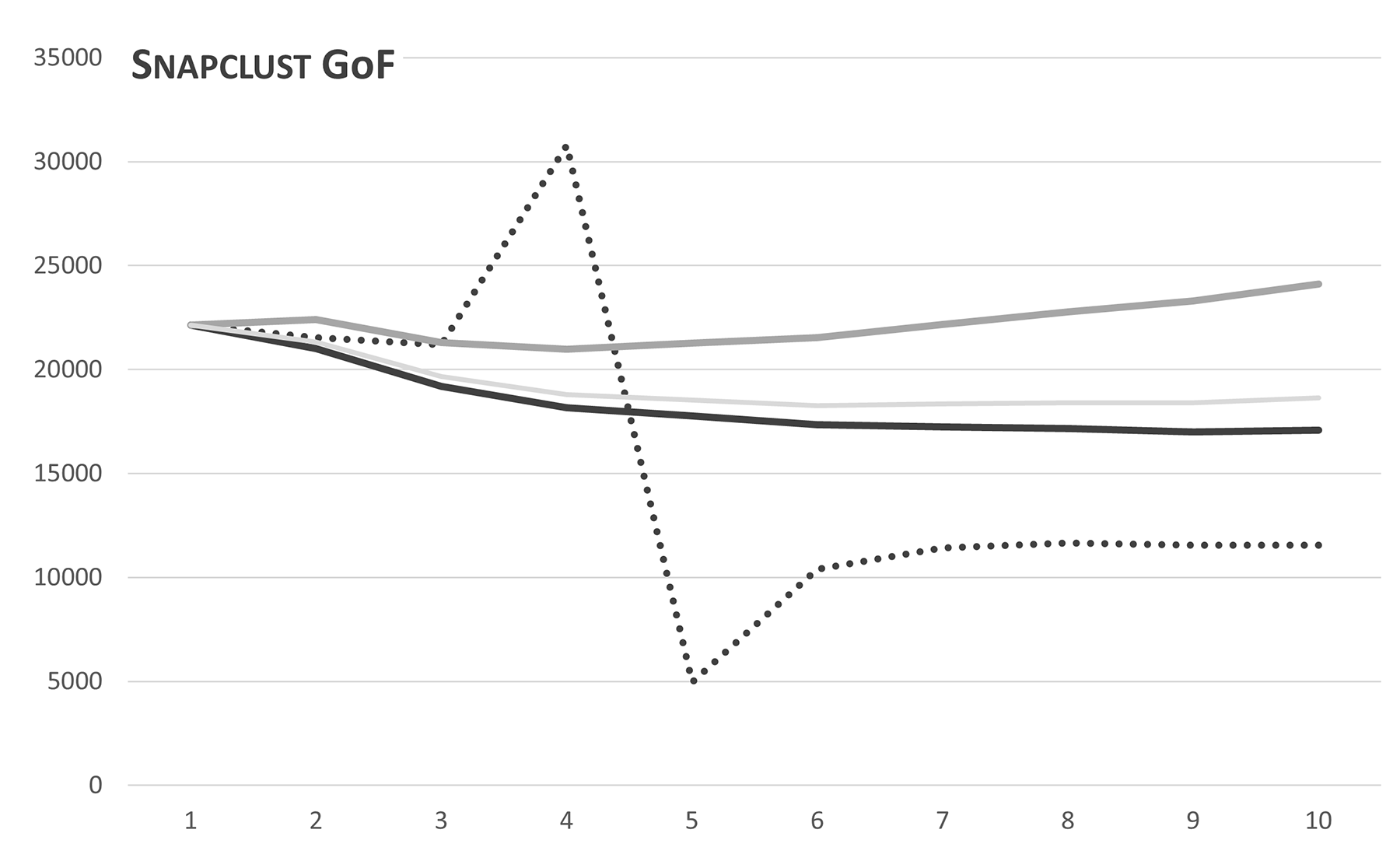

Each line corresponds to one of the goodness of fit statistic available with Snapclust (Jombart, 2008). The solid black line corresponds the AIC, the dotted black line to the AICc, the solid dark gray line to the BIC and the solid light gray line to the KIC.

### Supplementary Information 8. *Botrytis cinerea* strains and origin used to test host preference in microbiological tests.

| **Host plant for strain collection** | **Strain codes** | **Region** | **Location** | **Sampling date** | **Cultivar** |
| --- | --- | --- | --- | --- | --- |
| *Vitis vinifera*  (n=33) | VC200  VC217  VC224  VC228  VC271  VC280  VC288  VC297  VC302 | Champagne | Courteron  (n=9) | September 2006 | Pinot meunier |
|  | VC610  VC621  VC624  VC636  VC671  VC672  VC588 | Champagne | Hautvillers  (n=7) | September 2006 | Pinot noir |
|  | VC002  VC003  VC010  VC039  VC053  VC083  VC095 | Champagne | Vandières  (n=7) | September 2006 | Pinot noir |
|  | 14-V-300  14-V-306 | Alsace | Balbronn  (n=2) | June 2014 | Riesling |
|  | 14-V-007  14-V-019  14-V-022  14-V-028  14-V-029  14-V-063  14-V-220  14-V-221 | Alsace | Sigolsheim  (n=8) | June 2014 | Pinot auxerrois |
| *Solanum lycopersicum*  (n=11) | VC800  VC806  VC810  VC829  VC838  VC860 | Champagne | Courceroy  (n=6) | September 2006 | Moneymaker |
|  | VA708  VA714  VA721  VA734  VC902  VC911  VC927 | Champagne | Foissy-sur-Vanne  (n=7) | September 2005  September 2006 | Moneymaker |
| *Rubus fruticosus*  (n=7) | VC343  VC356  VC360  VC399 | Champagne | Courteron  (n=4) | September 2006 | Wild |
|  | VC141  VC150  VC172 | Champagne | Vandières  (n=3) | September 2006 | Wild |
| *Fragaria* x *ananassa*  (n=10) | 14-V-376  14-V-408  14-V-413  14-V-414 | Alsace | Balbronn  (n=4) | June 2014 | Cléry |
|  | 14-V-104  14-V-125  14-V-072  14-V-079  14-V-081  14-V-088 | Alsace | Sigolsheim  (n=6) | June 2014 | Candiss |
| *Hydrangea macrophylla*  (n=10) | 2252  2254  2293  2297  2334  2531  2556  2561  2668  2676 | Pays-de-Loire | Pont-de-Cé  (n=10) | ? | ? |
| Unknown (reference strain) | B05-10 | Lab. | - | - | - |
